# Pixel walking along the boreal forest–Arctic tundra ecotone: Large scale ground-truthing of satellite-derived greenness (NDVI)

**DOI:** 10.1101/2024.01.14.574721

**Authors:** Russell E. Wong, Logan T. Berner, Patrick F. Sullivan, Christopher S. Potter, Roman J. Dial

## Abstract

Satellite remote sensing of climate-driven changes in terrestrial ecosystems continues to improve, yet interpreting and rigorously validating these changes requires extensive ground-truthed data. Satellite measurements of vegetation indices, such as the Normalized Difference Vegetation Index (NDVI, or vegetation greenness), indicate widespread vegetation change in the Arctic that is associated with rapid warming. Plot-based studies have indicated greater vegetation greenness generally corresponds to greater plant biomass and deciduous shrub cover. However, the spatial scale of traditional plot-based sampling is much smaller than the resolution of most satellite imagery and thus does not fully describe how plant characteristics such as structure and taxonomic composition relate to satellite measurements of greenness. To improve interpretation of Landsat measurements of vegetation greenness in the Arctic, we developed and implemented a method that links satellite measurements with ground-based vegetation classifications. Here we describe data collected across the central Brooks Range of Alaska by field sampling hundreds of Landsat pixels per day, with a field campaign total of 23,213 pixels (30 m). Our example dataset shows that vegetation with the greatest Landsat greenness was taller than 1m, woody, and deciduous; vegetation with lower greenness tended to be shorter, evergreen, or non-woody. We also show that understory vegetation influences Landsat greenness. Our methods advance efforts to inform satellite data with ground-based vegetation observations using field samples at spatial scales more closely matched to the resolution of remotely sensed imagery.

## INTRODUCTION

Satellite-measured vegetation indices are critically important for documenting vegetation change in remote and inaccessible Arctic regions (Myneni et al., 1997; Berner et al., 2020; Myers-Smith et al., 2020). Since the 1970s, researchers have used vegetation indices such as the Normalized Difference Vegetation Index (NDVI), commonly referred to as “vegetation greenness” (hereafter greenness) as proxies for photosynthetic activity (Rouse et al., 1974; Running & Nemani, 1988; Beck et al., 2011; Berner & Goetz, 2022). Studies document increased greenness (greening) that is associated with warming in the Arctic (Bhatt et al., 2017; Berner et al., 2020) and along the northern edge of boreal forest (Beck et al., 2011; Berner & Goetz, 2022). Greening is generally interpreted as increased plant productivity due to strong correlations between greenness and field-based metrics of productivity such as leaf area index (LAI) and aboveground biomass (Sellers, 1985; Street et al., 2007; Berner et al., 2018).

Greenness has also been shown to vary with plant composition (Boelman et al., 2011; Anderson et al., 2016; Bendavid et al., 2023). For example, deciduous shrubs are generally “greener” than other plant functional types (PFTs), such as graminoids (Raynolds et al., 2008; Berner et al., 2018; Jespersen et al., 2023), thus suggesting that Arctic greening is partially driven by deciduous shrub expansion (Forbes et al., 2010; Myers-Smith et al., 2011; Fraser et al., 2014; Frost et al., 2014). Because greenness varies by plant type, many studies use greenness in combination with other metrics to predict and map landcover classes in the Arctic (Jorgenson et al., 2009; Macander et al., 2017; Walker et al., 2005). Other studies apply regression-based methods for predicting canopy cover using spectral endmember values of pixels containing pure landcover classes to estimate sub-pixel canopy cover of different Arctic plant types (Olthof & Fraser, 2007; Mikheeva et al., 2012; Nill et al., 2022).

Nevertheless, more ground data are needed to understand how plant characteristics such as LAI, biomass, height, and species composition at the landscape scale (10^2^-10^3^ km^2^) relate to satellite measurements of greenness (Myers-Smith et al., 2020; Berner & Goetz, 2022; Timoney, 2023). Meyers-Smith et al. (2020) highlighted two key issues in the spatial mismatch of satellite-derived greenness and field-based observations: poor replication of field observations across landscapes and the challenge in scaling-up field plot samples to image pixel size. Field studies that quantify vegetation characteristics including canopy cover, LAI, and biomass (Sellers, 1985; Street et al., 2007; Berner et al., 2018; Jespersen et al., 2023) are essential for understanding the biological determinants of greenness. However, due to their often-intensive sampling design, comprehensive plot-based field studies struggle logistically to characterize ecosystems with sufficient spatial continuity and coverage. While aggregates of ground-based data exist, they generally include datasets collected using different methods by different researchers over different time periods (Macander et al., 2022; Berner et al., 2024). This heterogeneity in sampling makes it difficult to accurately link satellite measurements with ground observations.

We are aware of no studies using consistent methods that have compared ground-truthed plant composition data to satellite-derived greenness at the scale of 1 km^2^ or more (i.e., ~10^3^ Landsat pixels of 30 m resolution). Here we illustrate a field method and its data analysis wherein we continuously sampled vegetation along 639 km of 30 m wide transects covering ~23,000 Landsat pixels. The method combines smartphone technology with extensive field sampling and prioritizes spatial scale over descriptive intensity. We have applied the field method, which we call “pixel walking,” during three seasons in Alaska’s Brooks Range, where we classified vegetation by PFT and cover along continuous ground-based field transects for up to 30 km per day and 60 days per field season. We then used these transects to characterize vegetation composition of individual Landsat pixels. To demonstrate the efficacy of pixel walking, we address three crucial questions linking Landsat imagery to ecosystems: First, can we use pixels of heterogeneous vegetation composition to characterize greenness of individual vegetation classes? Second, how does greenness differ among vegetation classes defined by PFT and structure? And third, how does vegetation structure (e.g., open versus closed canopy) affect greenness within individual vegetation classes?

Through analysis of an example dataset collected in 2020, we show both that greenness varies among PFTs such that the greenest plants are tall, woody, and deciduous, and that canopy structure affects greenness as perceived using Landsat imagery. We offer pixel walking as a novel approach that can improve not only the ecological interpretation of satellite-derived greenness, but also the ability to map vegetation composition across remote regions in the Arctic and elsewhere around the world.

## METHODS

To compare satellite-based NDVI to ground-based vegetation classifications over spatial scales >10^2^ km^2^, we collected vegetation data along field transects that sampled 10^2^-10^3^ Landsat pixels per day. During 22 July–17 August 2020, we recorded 639 km of vegetation transects near and across treeline in Gates of the Arctic National Park (mean elevation 704 m above sea level, range 300 – 1300 m asl), located within Alaska’s central Brooks Range (Fig. 1). Along each transect, we classified overstory and understory vegetation classes by PFT (Table 1). In all, we sampled 23,213 Landsat pixels, each with maximum 2020 greenness calculated using the *LandsatTS* R package (Berner et al., 2023). By extracting Landsat maximum greenness (hereafter “max-greenness”) values along the vegetation transects then applying a multiple regression model, we estimated for each vegetation class the mean of the max-greenness (“mean greenness” or *ndvi*).

**Figure 1.**
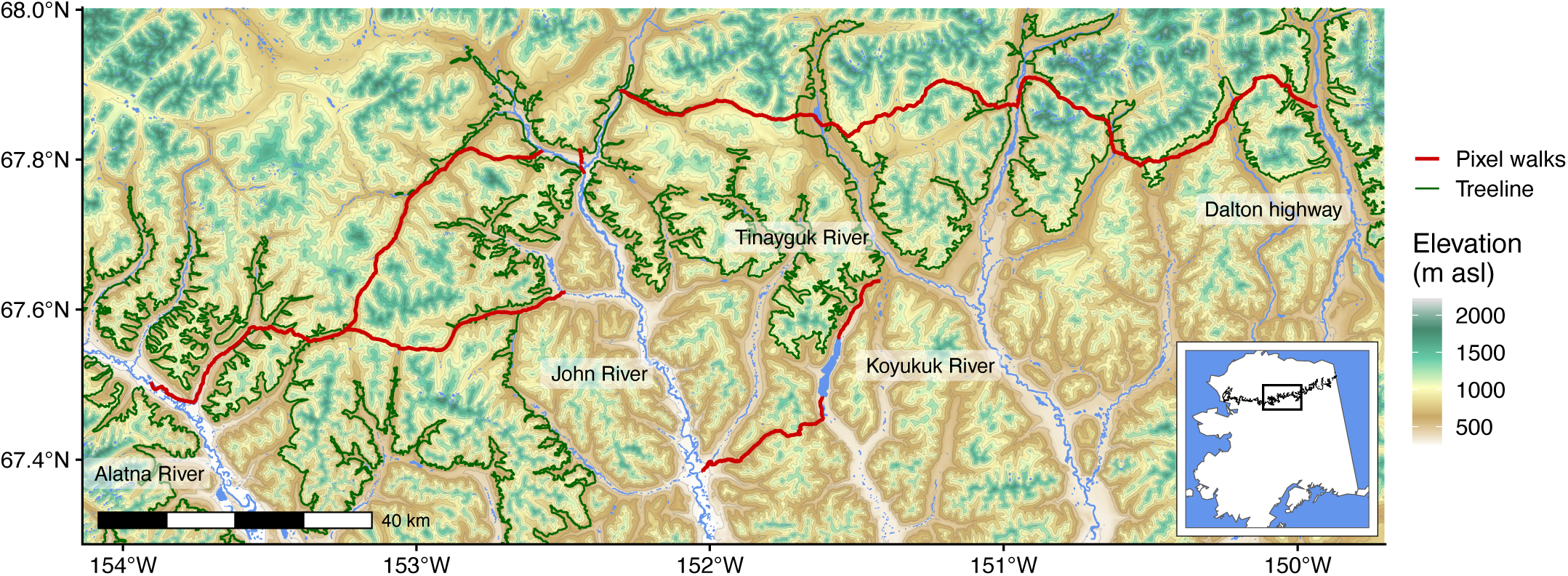
Study area and pixel walk transect locations. The transects started at the top right near the Dalton highway and continued counterclockwise ending on the Tinayguk River.

**Table 1.**
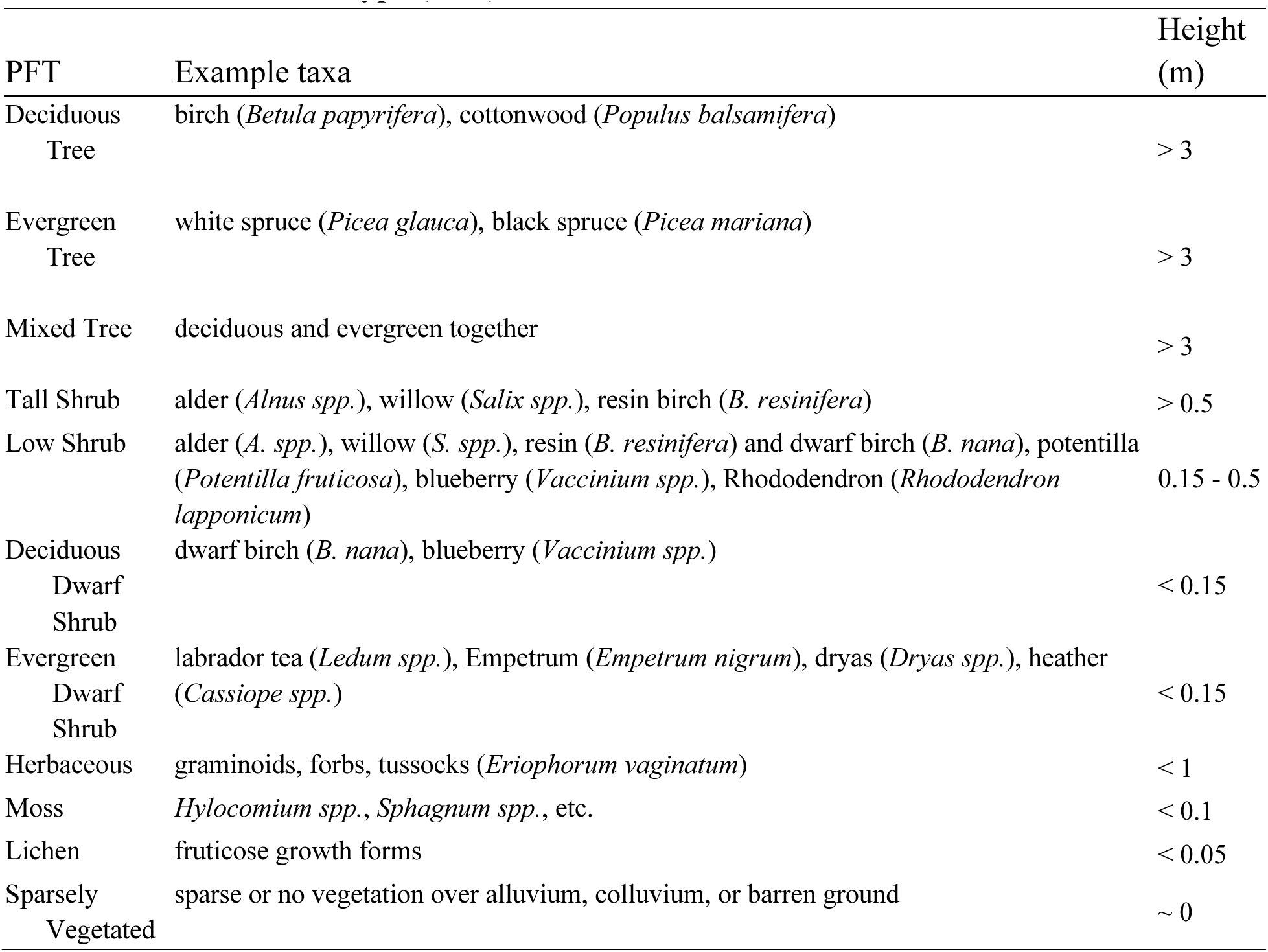
Plant functional type (PFT).

| PFT | Example taxa | Height (m) |
| --- | --- | --- |
| Deciduous Tree | birch (*Betula papyrifera*), cottonwood (*Populus balsamifera*) | > 3 |
| Evergreen Tree | white spruce (*Picea glauca*), black spruce (*Picea mariana*) | > 3 |
| Mixed Tree | deciduous and evergreen together | > 3 |
| Tall Shrub | alder (*Alnus spp.*), willow (*Salix spp.*), resin birch (*B. resinifera*) | > 0.5 |
| Low Shrub | alder (*A. spp.*), willow (*S. spp.*), resin (*B. resinifera*) and dwarf birch (*B. nana*), potentilla (*Potentilla fruticosa*), blueberry (*Vaccinium spp.*), Rhododendron (*Rhododendron lapponicum*) | 0.15 - 0.5 |
| Deciduous Dwarf Shrub | dwarf birch (*B. nana*), blueberry (*Vaccinium spp.*) | < 0.15 |
| Evergreen Dwarf Shrub | labrador tea (*Ledum spp.*), Empetrum (*Empetrum nigrum*), dryas (*Dryas spp.*), heather (*Cassiope spp.*) | < 0.15 |
| Herbaceous | graminoids, forbs, tussocks (*Eriophorum vaginatum*) | < 1 |
| Moss | *Hylocomium spp.*, *Sphagnum spp.*, etc. | < 0.1 |
| Lichen | fruticose growth forms | < 0.05 |
| Sparsely Vegetated | sparse or no vegetation over alluvium, colluvium, or barren ground | ~ 0 |

### Vegetation classification

We classified vegetation composition using a modification of the Alaska Vegetation Classification system (Viereck et al., 1992) that described the two uppermost strata: overstory and understory. Within each stratum, we recorded PFT class as one of 11 classifications (Table 1) and canopy cover as one of two classifications (25% < open < 50% < closed < 100% cover). When PFT canopy cover was less than 25%, that PFT was not recorded. Such areas with low PFT cover (< 25%) above barren ground or alluvium were classified as “sparsely vegetated.”

### Pixel walking transects

Using Gaia GPS (gaiagps.com) on a recent-model iPhone, we recorded daily GNSS tracks of 40 individual field transects, each 1–28 km long (Fig. 1). Each transect was one continuous GNSS track of vegetation recorded by a single individual on a single day. Two observers recorded side-by-side transects simultaneously while walking single file. Because we traveled most efficiently along vegetation edges (e.g., boundaries between tall shrubs and lower statured PFTs), each observer considered the vegetation on a fixed side of the directed GNSS track. One observer sampled to the left while the other to the right. To match Landsat pixel resolution (30 m), we considered only patches of vegetation sized ~900 m^2^ or larger, constrained by location within 30 m perpendicular to the line of travel. If the vegetation at the 900 m^2^ scale changed in PFT or canopy cover, we located and documented the “changepoint” (Fig. S1 and Supplementary Methods) on our iPhones using a form-based data collection app (e.g., FormConnect Pro, formconnections.com). Changepoints recorded the location, date, time, side (left or right), PFT for each stratum, and observer.

### GPS track attributing

We used R (4.3.1) and QGIS (3.32.3) to re-draw each day’s directed walk as two directed paths, one located 15 m to the left and another 15 m to the right of the directed walk to represent the center of adjacent 30 m pixels (Fig. S1 and Supplementary Methods). Because the directed walks were GNSS tracks, both the GNSS track and the two directed paths 15 m away consisted of segments separated by vertices. For each changepoint *i* we located the nearest path vertex to the left or right and attributed that vertex and every forward vertex up to the next changepoint *i + 1* with the attributes of changepoint *i*. These vertices, together with their attributes, were used to create spatial line dataframes in the *terra* package (v. 1.7-55; Hijmans et al., 2023) in R.

### Landsat pixel processing

For comparison to the landscape surrounding the pixel walks but within the geographic extent of the directed paths, we extracted the max-greenness pixels intersected by the directed paths and for 20,000 random off-path pixels. To calculate and extract max-greenness values from pixels that were intersected by spatial lines representing directed paths, we found the pixel centers along the directed paths and exported Landsat data for them in Google Earth Engine (Gorelick et al., 2017) using the *LandsatTS* R package (Berner et al., 2023). We similarly exported Landsat data for the random landscape samples. Specifically, we exported surface reflectance measurements from Collection 2 of Landsat 5, 7, and 8 images. We filtered measurements by removing those with water, snow, cloud cover > 80%, geometric uncertainty > 15 m, and solar zenith angle > 60°. After filtering, 8-9 observations per pixel remained over the 2020 growing season (31 May to 31 August). We then calculated NDVI, cross-calibrated the 2020 Landsat 5 and 8 NDVI values with Landsat 7 using a machine learning approach, and finally estimated max-greenness using phenological modeling, all in *LandsatTS*.

### Estimating greenness values

Because individual pixels are often heterogeneous, the goal of our analysis was to determine a mean greenness value (*ndvi*) for each vegetation class defined as plant functional type (PFT), stratum, and–for overstory PFTs only–the cover class (open or closed). For example, some pixels contained both open- and closed-canopy PFTs, and so we estimated *ndvi* for each combination of canopy cover and PFT (covrPFT). Furthermore, because understory vegetation may contribute to greenness values when viewed from above, we separated the *ndvi* of PFTs in two ways. One way was as closed-canopy PFTs (cPFT) or open-canopy PFTs (oPFT). The second way was as a combination of covrPFT with understory PFT.

Of the 23,213 pixels sampled slightly more than half were homogeneous in cPFT. Of the total, 20,335 had at least one cPFT and 7,126 had at least one oPFT. Mean greenness of cPFTs was estimated with the 20,335 pixels that had cPFTs, of which 13,790 (68%) had only a single cPFT. Mean greenness of oPFTs was estimated with the 7,126 pixels that had oPFTs, of which 5,438 (76%) had only a single oPFT.

Here we present mean greenness estimated both as the arithmetic mean of pixels with a single PFT and as coefficients from a multiple regression model that incorporates sub-pixel variability in PFTs, thereby retaining over 8,000 pixels for analysis. To estimate *ndvi* of each covrPFT, we used multiple regression without intercept, where the dependent variable was pixel NDVI, the independent variables were the proportion of the pixel walk in that pixel of a given covrPFT, and samples were pixels crossed by pixel walks. This approach is similar to that used by Jespersen et al. (2023; Supplementary Text).

### Multiple regression unmixing

For covrPFT*_i_* in pixel-*j*, we determined the length of the spatial line within pixel-*j* and the length of the line with covrPFT*_i_*, then calculated the proportion, *p_ij_,* as the length of the line with covrPFT*_i_* inside pixel-*j* divided by the total line length in pixel-*j* (Supplementary Methods). We assumed that max-greenness in pixel-*j* (NDVI*_j_*) within a Landsat pixel was a linear combination of the *ndvi* values across all covrPFTs within that pixel. Thus, the max-greenness of pixel-*j* (*NDVI_j_*) can be represented as the sum (over the total *n* covrPFTs) of greenness (*ndvi_i_*) of each covrPFT_i_ weighted by the proportion (*p_ij_*) of pixel-*j* covered by PFT*_i_*:

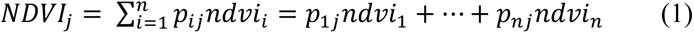

To solve for the unknown *ndvi_i_* across all pixels traversed, we implemented (1) using the function **lm()** in R as a multiple regression model without intercept:

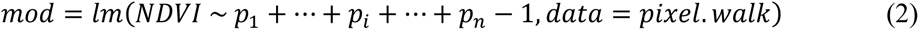

In (2), *pixel.walk* is a data frame where each row is a pixel and each of the *n* columns, *p_1,_ p_2,_ p_3,…_ p_n_*, is a covrPFT with variables representing the proportions of each covrPFT. The coefficients and standard errors associated with each *p_i_* as found using **summary(mod)** in R provided estimates of *ndvi_i_* mean greenness values.

While we included all covrPFTs that we encountered on the pixel walks across all 23,213 pixels (i.e., *pixel.walk* had 23,213 rows) in the multiple regression, we report only the greenness values of covrPFTs with a sample length greater than 1 km. Because n = 30 pixels represents about 1 km, we considered samples of less than 1 km (approximately 30 pixels) as unreliable for estimating *ndvi*. We considered the *ndvi* of any pair of covrPFT as significantly different at the 0.05 significance level, if their mean ± 2se did not overlap. We repeated the regression for all combinations of covrPFT and understory PFT, but only report the estimates where combinations of overstory and understory PFT were observed with > 1km of open and closed canopy.

### Arithmetic pixel means from homogeneous pixel samples

As a check against the ndvi_i_ estimates found using (2), we used those pixels with homogeneous covrPFT cover to calculate mean greenness ndvi_i_ values for each covrPFT. To calculate arithmetic pixel means for each covrPFT, we took the sum of the NDVI for each single-covrPFT pixel and divided by the sample size. Arithmetic pixel means are reported only for those covrPFTs with a sample length greater than 1 km. Approximately 59% of all sampled pixels were characterized by one cPFT: 28% had a mix of one cPFT and at least one other covrPFT; and 12% had only oPFTs. Approximately 24% of all sampled pixels were characterized by one oPFT; 6% had a mix of one oPFT and at least one other covrPFT; and 87% had only cPFTs.

## RESULTS

We traversed 23,213 total Landsat pixels during 608 km of pixel walks, sampling 11 cPFTs and five oPFTs *en route*. We report the arithmetic means from 13,790 single-cPFT pixels and regression means from 20,335 pixels with one or more cPFTs. We also report arithmetic means from 5,438 pixels of single-oPFTs and regression means from 7,126 pixels with one or more oPFTs.

Over half (59.5%) of the pixel walks passed through shrub dominated cPFTs (362 km), 30% through trees (224 km), and most of the remainder through herbaceous (20 km, 3%). Overall, the mean greenness across all pixel walks (mean = 0.65, sd = 0.13, n = 23,213 pixels) was very similar to the landscape (mean = 0.67, sd = 0.15, n = 19,550 pixels); however, the distribution we sampled was slightly less green than the landscape overall (Fig. 2).

**Figure 2.**
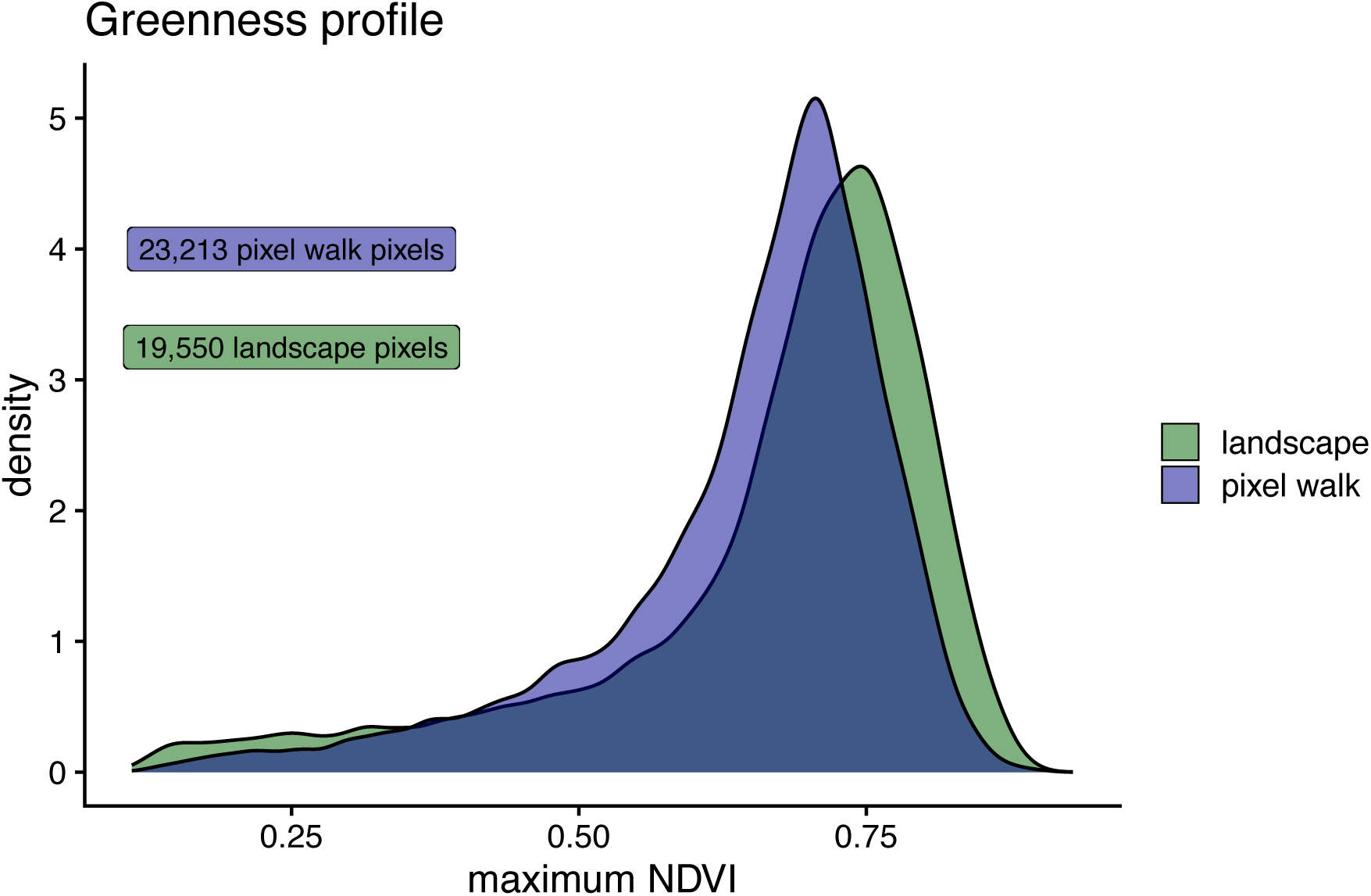
Density profile of maximum NDVI (greenness) across two samples. The pixel walk (blue) is all pixels sampled using vegetation transects. The landscape (green) is a random sample of all pixels within the spatial extent of the pixel walk but excluding those pixels sampled using vegetation transects. Our goal was to describe and compare mean greenness among PFTs, rather than use pixel walks to estimate landscape mean greenness.

### Homogeneous vs. heterogeneous pixel estimates of PFT greenness

Arithmetic pixel means demonstrated similar mean values to regression coefficient estimates (Fig. 3 and S2) but were slightly lower, except for evergreen and deciduous trees which were slightly larger. Regression of all pixels, regression of single-PFT pixels, and arithmetic pixel means were not significantly different with differences generally less than 0.004 NDVI units on average. Variability, measured as two standard errors, was also similar.

**Figure 3.**
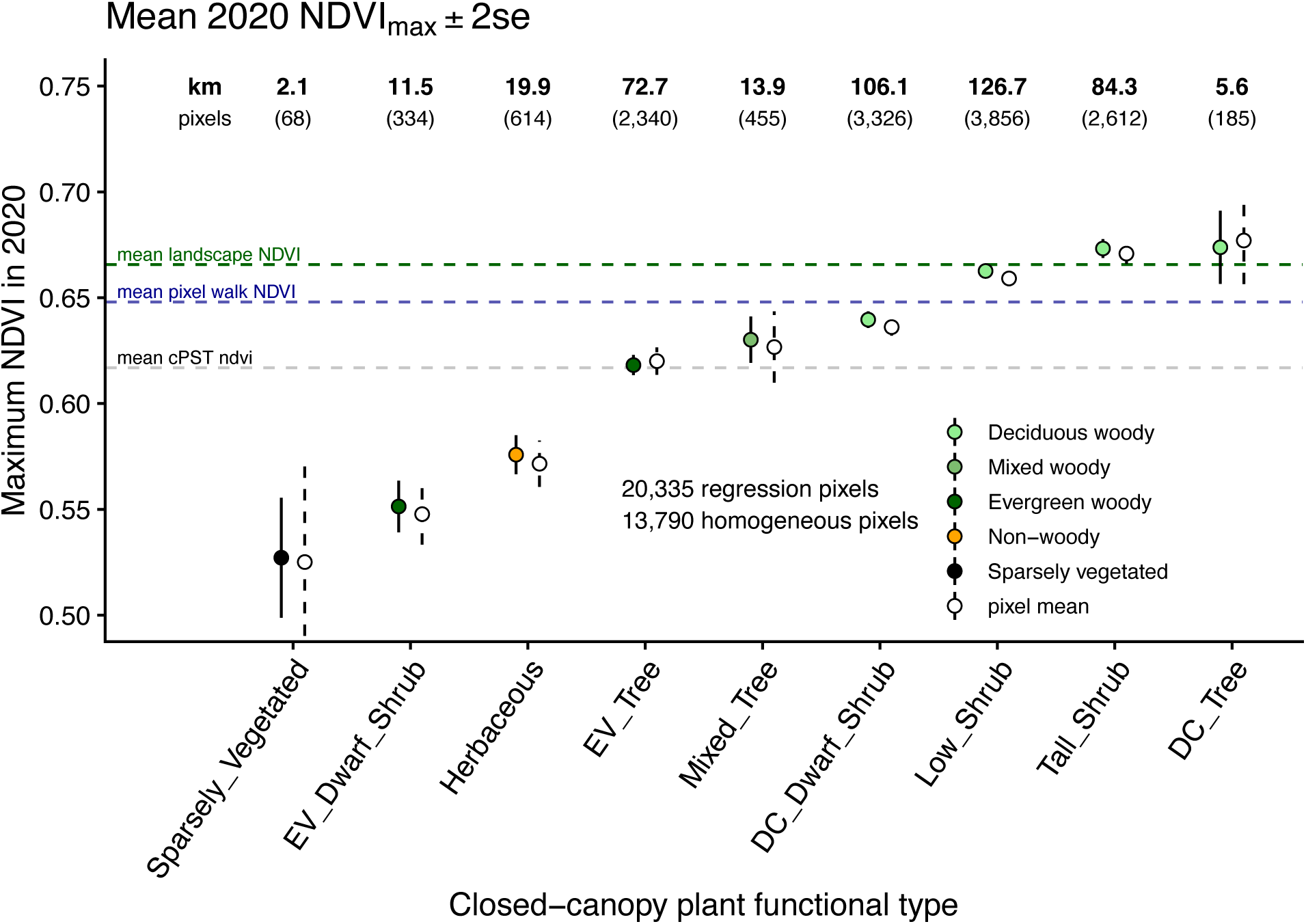
Greenness values for each closed-canopy plant functional type (cPFT). Filled circles represent multiple regression greenness values and colors represent coarse vegetation classes. Open circles are the arithmetic pixel means calculated across all single-cPFT pixels. Error bars represent two standard errors. Above numbers represent km of each cPFT sampled (bold) for multiple regression and number of pixels (parentheses) used to calculate the arithmetic pixel means. The horizontal dashed lines are the mean NDVI of ~20,000 pixels outside of the pixel walk (green), mean NDVI of ~20,000 pixels sampled by the pixel walk (blue), and mean NDVI of all nine PFTs. Deciduous (“DC”) and evergreen (“EV”) are abbreviated in the x-axis labels.

### Closed canopy PFTs

Among all cPFTs, those with deciduous vegetation were the greenest on average (mean = 0.66, n = 4 cPFTs), surpassing the overall mean of 0.62 averaged across all nine cPFTs (Fig. 3). The least green was evergreen dwarf shrub (0.55 ± 0.012 [mean ± 2 SE], d = 11.5 km). Surprisingly, greenness of the sparsely vegetated cPFT (0.53 ± 0.028, d = 2.1 km), while possibly under-sampled and highly variable in reflectance, was not significantly different from evergreen dwarf shrubs.

Among the four shrub cPFTs, taller cPFTs were greener than shorter ones (Fig. 3). Of the shrub cPFTs, tall shrub (0.67 ± 0.004, d = 84.3 km) and low shrub (0.66 ± 0.004, d = 126.7 km) shared above average greenness estimates, and evergreen dwarf shrub (0.55 ± 0.012, d = 11.5 km) displayed the lowest greenness estimate of all cPFTs (Fig. 3).

Greenness of closed deciduous trees (0.67 ± 0.018, d = 5.6 km) was greater than greenness of closed evergreen trees (0.62 ± 0.004, d = 72.7 km; Fig. 3). Greenness of closed mixed trees (0.63 mean ± 0.010, d = 13.9 km) was intermediate between closed deciduous and evergreen trees but only slightly greater than closed evergreen trees. Greenness of closed deciduous trees was greater than the overall mean cPFT greenness (0.67 > 0.62) while the lower and upper bounds of two standard errors of closed mixed trees and closed evergreen trees, respectively, overlapped with the overall mean cPFT greenness.

### Open vs. closed canopy PFTs

In general, greenness of open-canopy shrubs was similar to greenness of their closed-canopy counterparts (Fig. S2). Closed deciduous tall shrubs (0.67 ± 0.004, d = 84.3 km) were not significantly different than open deciduous tall shrubs (0.69 ± 0.022, d = 3.4 km). Open low shrubs (0.66 ± 0.004, d = 23.3 km) and closed low shrubs (0.66 ± 0.004, d = 126.7 km) were nearly identical. The exception was that closed deciduous dwarf shrubs (0.64 ± 0.004, d = 106.1 km) were significantly greener than open deciduous dwarf shrubs (0.51 ± 0.016, d = 6.4 km).

Greenness of open-canopy trees differed significantly from greenness of closed-canopy trees (Fig. S2). For example, open deciduous trees (0.58 mean ± 0.032, d = 1.6 km) were significantly less green than closed deciduous trees (0.67 mean ± 0.018, d = 5.6 km). Conversely, open evergreen trees (0.67 mean ± 0.004) were greener than closed evergreen trees (0.62 mean ± 0.004), a difference explored below. Notably, open evergreen trees (d = 130.5 km) were sampled twice as much as closed evergreen trees (d = 72.7 km) and more than any other covrPFT; however, both open and closed deciduous trees had relatively small sample lengths.

The understory contributed to the greenness of pixels. For example, deciduous understories of open PFTs were generally greener than non-deciduous understories of the closed PFT counterpart (Fig. 4). An exception was low shrub over evergreen dwarf shrub. The open canopy version was greener than the closed canopy version even though the understory was not deciduous. Notably, evergreen trees were generally greener as open canopy, because their understories were mostly tall and low deciduous shrubs (Supplementary Text).

**Figure 4.**
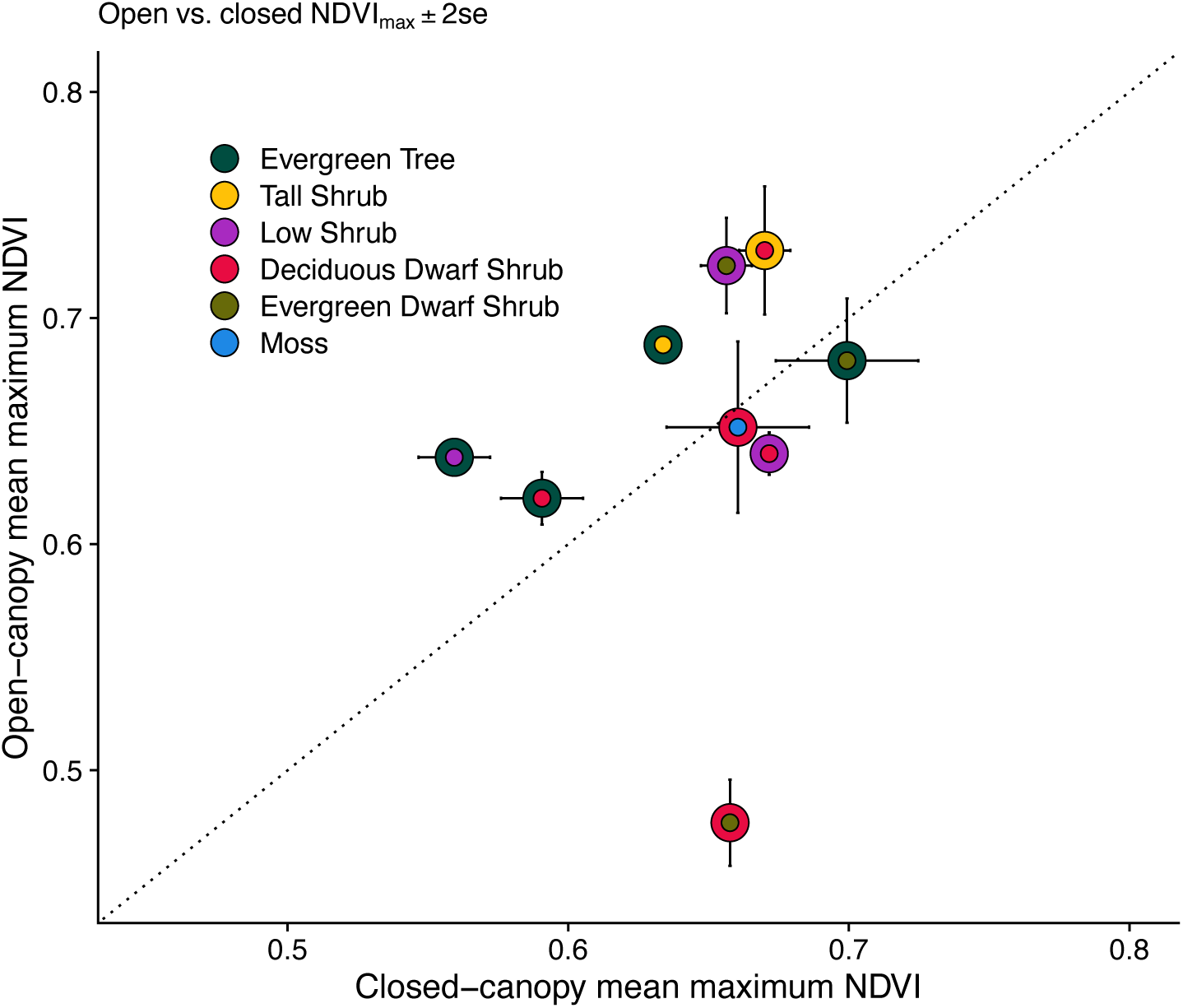
Greenness values for closed and open canopy overstories with different understories. Outer circles represent overstory PFT, and inner circles represent understory PFT. Only overstory PFTs had canopy cover classification. The line of equality is dotted.

## DISCUSSION

We have demonstrated a novel, large-scale approach to ground truthing that informs the relationship between Landsat-derived greenness and ground-based vegetation. By prioritizing spatial scale over detail of ground-based data, we characterized the vegetation of 23,213 Landsat pixels using 639 km of vegetation transects. We used the greenness of Landsat pixels to estimate greenness values of individual PFTs with two methods: multiple regression of all pixels and arithmetic means of single-PFT pixels, which yielded similar results.

We found that PFTs with greater greenness were tall, woody, and deciduous. PFTs with lower greenness were short, non-woody, or evergreen. Furthermore, understory vegetation influenced satellite greenness measurements of all PFTs and, within individual PFTs, accounted for greenness differences between open versus closed canopies. From a satellite perspective, greenness depends on the combination of overstory and understory vegetation.

### Landscape greenness

In Alaska, treeline is advancing in some places (Dial et al., 2022) but not others (Rees et al., 2020). Similarly, greening trends suggest a boreal forest biome shift (Beck et al., 2011; Berner & Goetz, 2022). However, simulations suggest that forest expansion may not always lead to greening (Huemmrich et al., 2021). Given their relatively low greenness, evergreen tree expansion and infilling may only be detectable as greening when trees are growing over evergreen dwarf shrubs, herbs, or sparse vegetation. On the other hand, canopy closure of evergreen trees overtopping tall deciduous shrubs could result in browning, a decrease in NDVI over time (Huemmrich et al., 2021).

The overall greenness of the landscape suggests a prevalence of low to tall shrubs within our study area, consistent with our expectations and regional vegetation maps (Walker et al., 2005; Jorgenson et al., 2009; Macander et al., 2017). However, the difference between landscape greenness and transect greenness suggests a sampling bias towards less green vegetation, which are often shorter and easier to walk through. While our pixel walk did not provide a representative sample of the landscape, it nevertheless enabled us to collect detailed ground data about the full range of vegetation composition and structure present on the landscape. These data then allowed us to rigorously investigate how vegetation structure and composition influence Landsat greenness values.

### Closed PFTs (cPFTs)

We found the greenest cPFTs on average were deciduous trees and shrubs, and that woody plants were generally greener than non-woody plants. These findings are consistent with prior plot-based studies from the Arctic (Raynolds et al., 2008; Yang et al., 2019; Spadoni et al., 2020; Jespersen et al., 2023). Moreover, we found taller woody plants are greener than shorter woody plants, likely because they have greater LAI. Our observation that tall deciduous shrubs are typically greener than all other tundra PFTs supports the interpretation that Arctic greening is linked to expansion and increasing biomass of tall deciduous shrubs over low-lying tundra, especially graminoids and lichens (Forbes et al., 2010; Myers-Smith et al., 2011; Fraser et al., 2014; Frost et al., 2014).

### Open vs closed canopy PFTs

Satellite measurements of greenness are affected by both overstory and understory vegetation, which can complicate interpretations of sub-pixel greenness. For example, 90% of open evergreen tree samples had an understory of deciduous shrubs, increasing the greenness of open evergreen trees relative to closed evergreen trees. Similarly, other studies have shown that forest greenness is positively correlated with understory shrub cover, while weakly or negatively correlated with overstory conifer cover due to masking understory shrubs (Loranty et al., 2018; Bendavid et al., 2023).

In contrast to evergreen trees, open deciduous dwarf shrubs were less green than closed deciduous dwarf shrubs. This difference likely resulted because both open and closed deciduous dwarf shrubs usually grew over less green PFTs, such as evergreen dwarf shrubs and lichen (Fig. 4 and Supplementary Text). Open tall and low shrubs were similar in greenness to closed tall and low shrubs, probably because these communities were mostly composed of deciduous overstories and understories. Alternatively, satellite measurements of greenness may begin to saturate in areas with high shrub biomass (Raynolds et al., 2012; Berner et al., 2018; Bonney et al., 2018), which may explain the small differences in greenness between open and closed tall and low shrubs. Therefore, the steepest compositional greening trends may be associated with range expansion of deciduous shrubs, with shrub infilling accounting for slower greening trends. This is an area in need of future study.

### Broader applications of pixel walking

Pixel walking is an approach for spatially extensive ground data collection that can be used for a variety of remote sensing applications. Here, we demonstrated how pixel walking can provide ground data on vegetation composition and structure to enable better understanding of Landsat greenness measurements in Arctic tundra and, by extension, greening trends across these northern lands. Nevertheless, this approach could also be used, for instance, to collect ground data for mapping vegetation cover, health, and disturbance impacts.

### Conclusions

Pixel walking is a novel and informative method that can provide data at an intermediate scale between satellite imagery and fine-scale, plot-based measurements. Our analysis showed Landsat-derived greenness varies among PFTs in previously poorly understood ways, such as how understory greenness influences satellite-observed greenness, and that satellite-observed greenness is positively related to shrub height. Further development of pixel walking could include analyses of vegetation composition and greening as well as more detailed classifications.

The technology and methods for satellite remote sensing continue to improve our ability to document vegetation change across the globe. However, much of the sub-pixel information about these vegetation changes remains obscured, given the resolution of Landsat sensors. Our methods advance efforts to link satellite imagery with ground-based vegetation by collecting data at spatial scales that better represent the capabilities of modern remote sensing.

## ACKNOWLEDGEMENTS

R.E.W was supported by National Aeronautics and Space Administration (NASA) through the Alaska Space Grant Program under grant no. 80NSSC20M0070. L.T.B. was supported by the NASA’s Arctic Boreal Vulnerability Experiment (ABoVE) under grant no. 80NSSC22K1247. P.F.S. was supported by National Science Foundation (NSF) awards OPP-1748849 and DEB-2133494. R.J.D was supported by NSF award OPP-1748773, Alaska NASA EPSCoR, an Explorers Club/Discovery Grant, and Shoreline, Incorporated. We appreciate the assistance in field data collection by A. Dahl, J. Ditto, S. Donahue, R. Koleser, K. Martin, T. Matsuoka, S. Smeltz, K. Vicich, and M. Zietlow.

## Supplementary Figures

**Figure S1.**
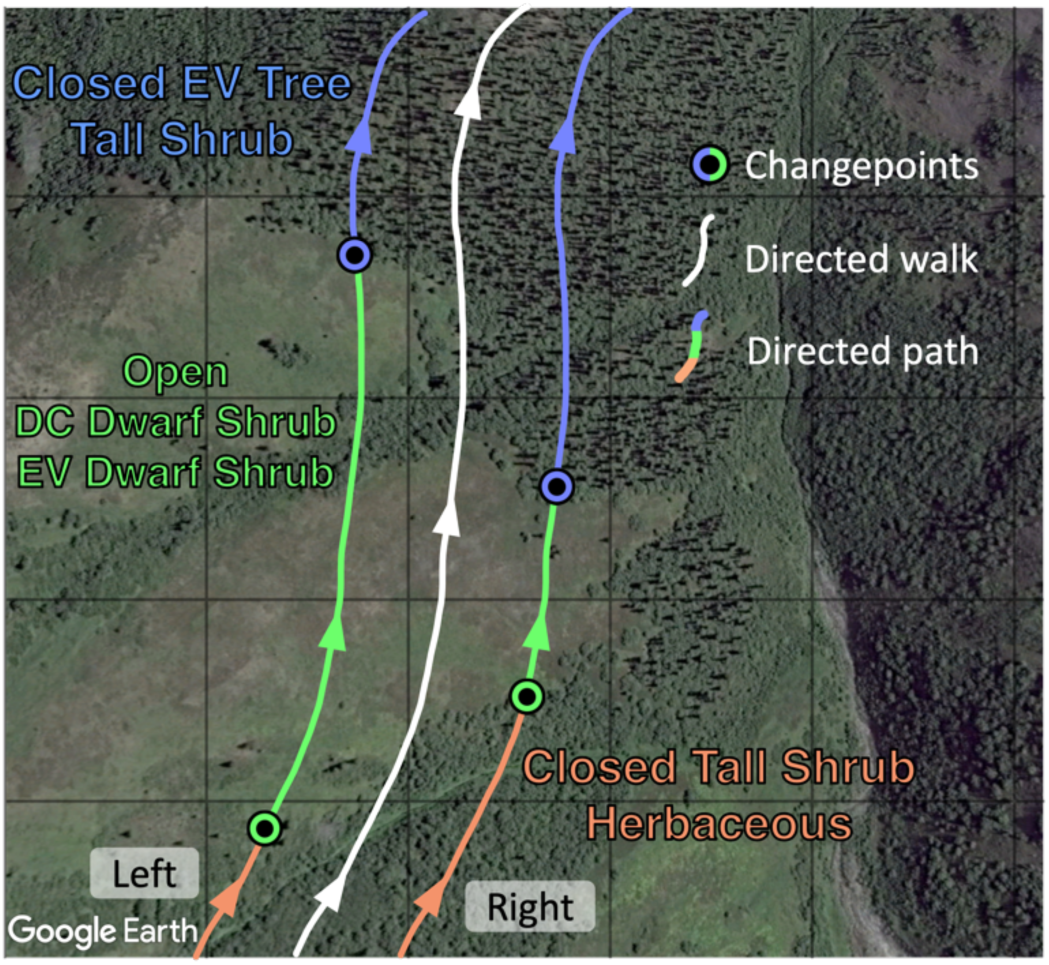
Conceptual illustration of the directed walk (white) with left and right directed paths (colored) on either side. The directed walk is the actual line of travel in the field. Out of the field, the directed paths were generated from the directed walk GNSS track and represent the vegetation observed by two field technicians that classified vegetation looking to either the left or right of the walk. The directed paths are 15 m away from the directed walk and 30 m away from each other. Each changepoint records where the vegetation changes and designates the vegetation class along directed path until another changepoint is collected. The grid represents Landsat pixel scale (30 m).

**Figure S2.**
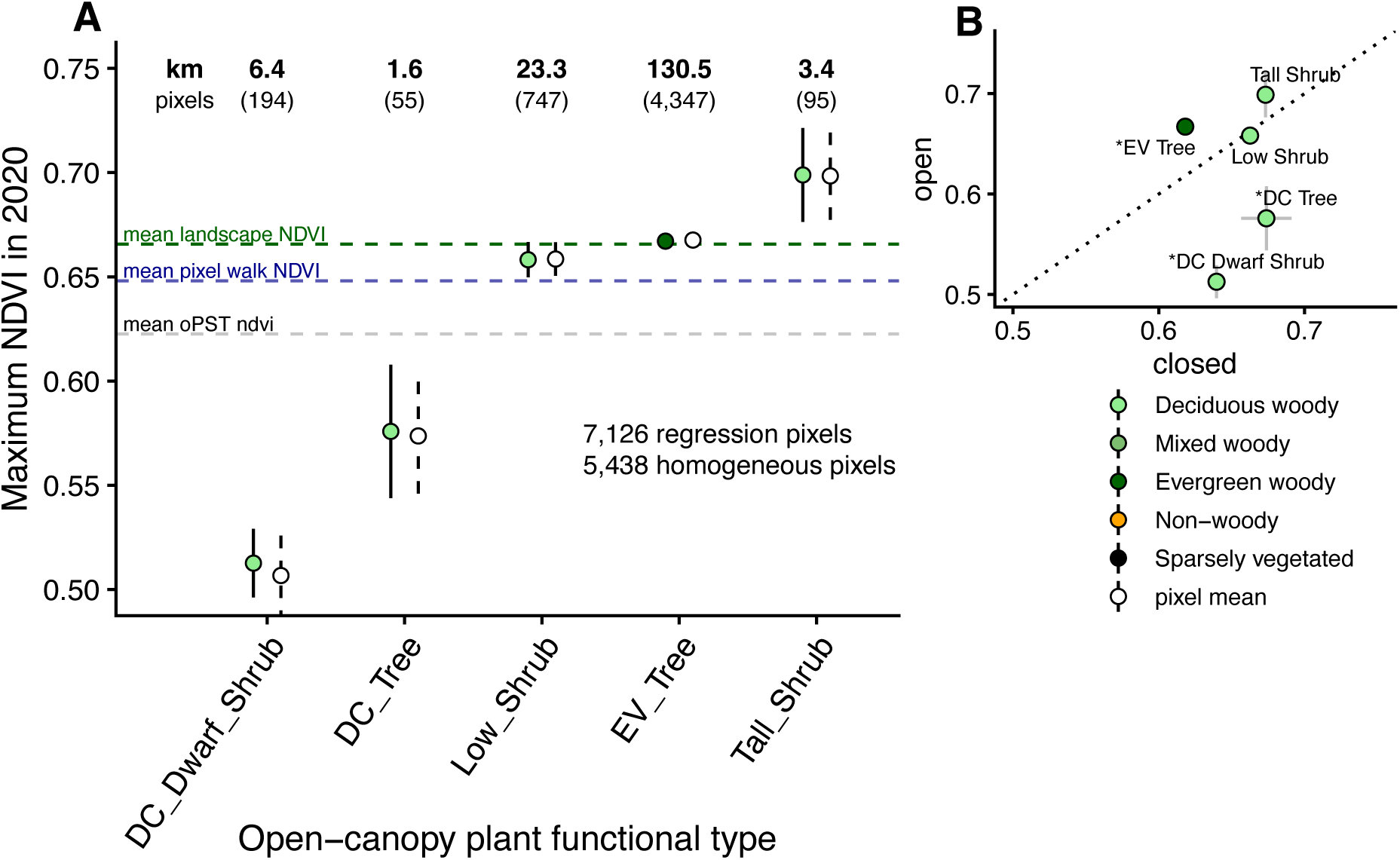
**(A)** Greenness of open-canopy PFTs (oPFTs). Filled circles represent multiple regression greenness values and colors represent coarse vegetation classes. Open circles are the arithmetic pixel means calculated across all single-oPFT pixels. Error bars represent two standard errors. Numbers in bold at top of figure represent sample sizes in kilometers and numbers in parentheses sample sizes in pixels. The horizontal dashed lines are the mean NDVI outside of the pixel walk (green), mean NDVI of ~20,000 pixels sampled by the pixel walk (blue), and mean NDVI of all nine PFTs. **(B)** Open-vs. closed-canopy ndvi for five PFTs. Line of equality is dotted. The asterisk (*) indicates where two standard errors do not cross line of equality.

## Supplementary Methods

### GPS track attributing

Here we demonstrate the methods for processing a pixel walk dataset. The process includes re-drawing and attributing each day’s directed walk as two directed paths, one 15 m to the left and one 15 m the right (Fig. Sxx). The following is a demonstration of this process for the directed *path* to the left of one day’s directed *walk.* Most of the process is completed in R, with a manual step completed in QGIS. Here are the steps:

- bring in directed walk as a raw GPS track
- create points along a 15 m buffer around the track
- manually choose the side of the buffer that will be used as the re-drawn GPS track (directed path)
- order the points from start to end
- attribute ordered points with changepoint values
- convert ordered points to spatial lines

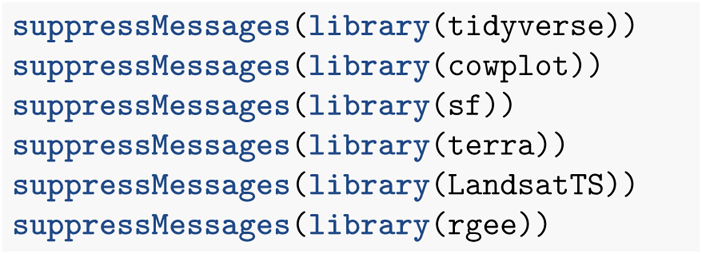

### Bring in directed walk as a raw GPS track

First, we brought in the raw GPS track. This track was recorded on Gaia GPS, then exported as a KMZ. Because **sf::st_read()** does not read KMZs, we used Google Earth Pro to convert the KMZ to KML.

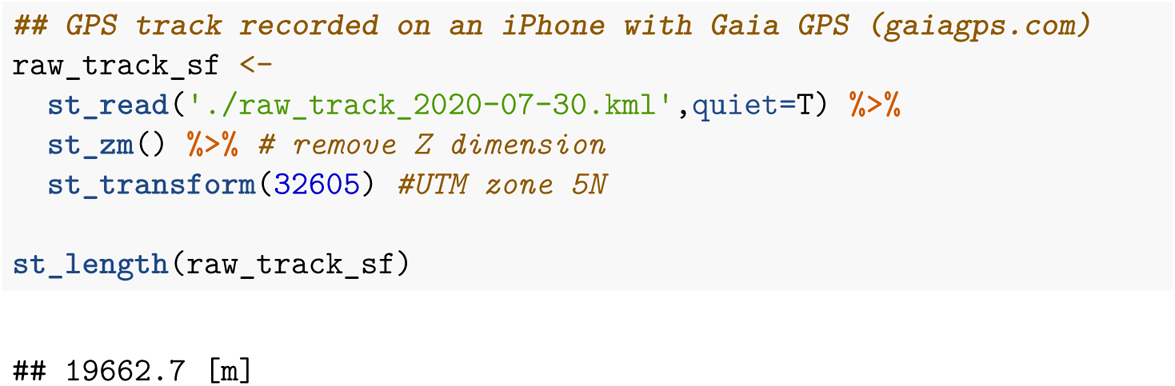

### Create points along a 15 m buffer around the track

We created a buffer 15 m away from the raw GPS track using **sf::st_buffer()**. We then converted the buffer (polygon) into lines using **sf::st_cast(“LINESTRING”)** and sampled one point every 5 m along the line using **sf::st_line_sample()**. These buffered points would become the vertices of the directed paths 15 m to the left and right of the raw GPS track.

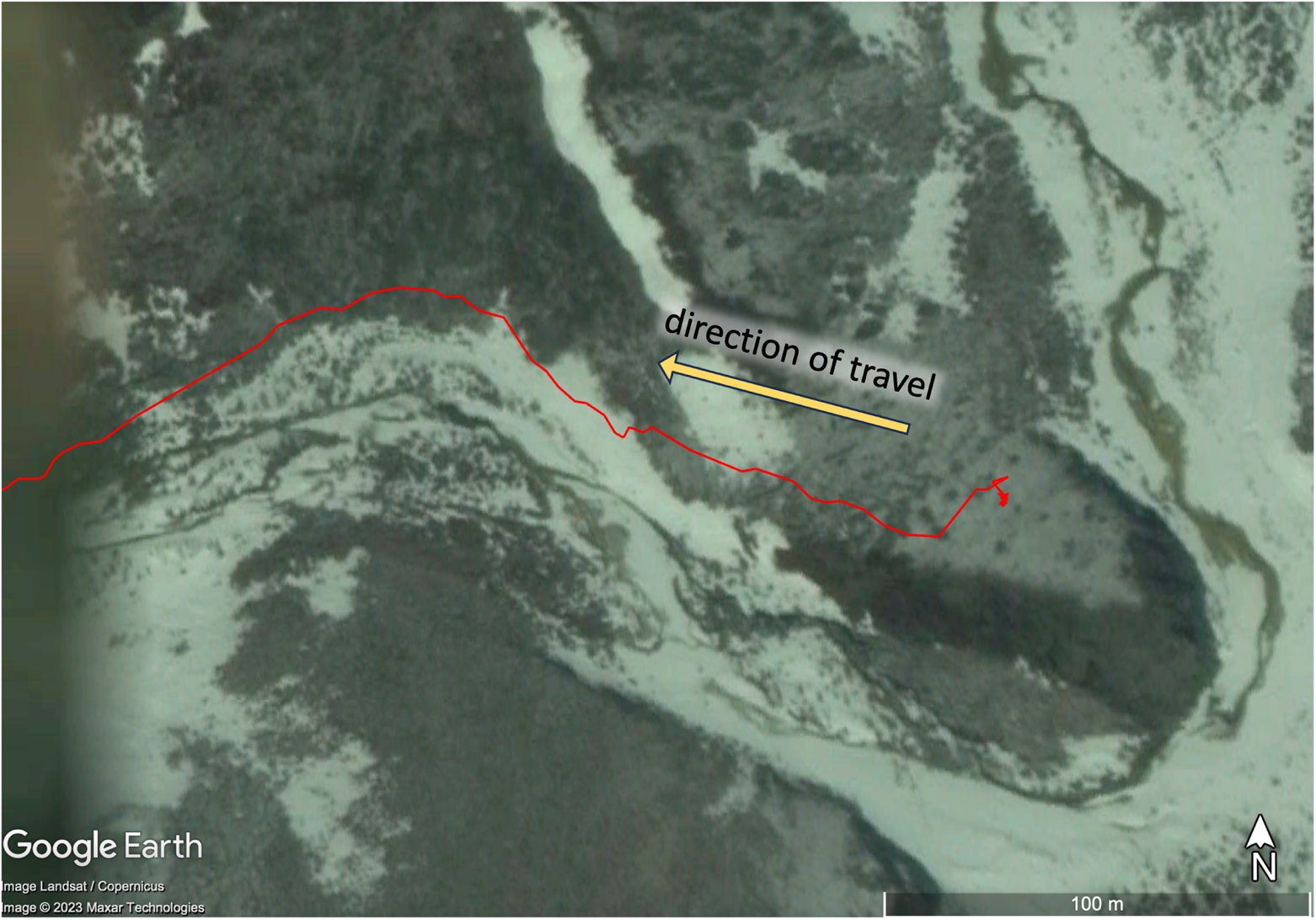
A portion of a directed walk recorded as a raw GPS track (red) collected on July 30, 2020. The yellow arrow shows the direction of travel.

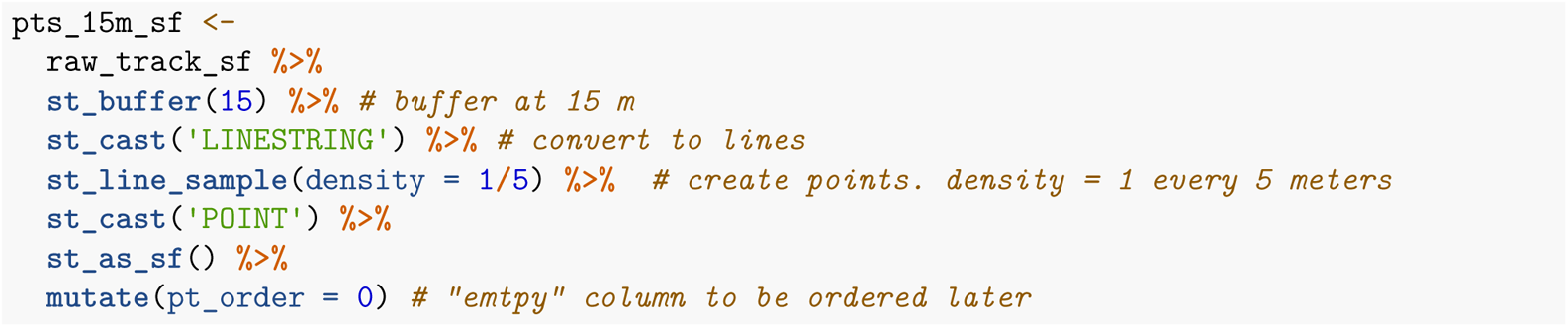

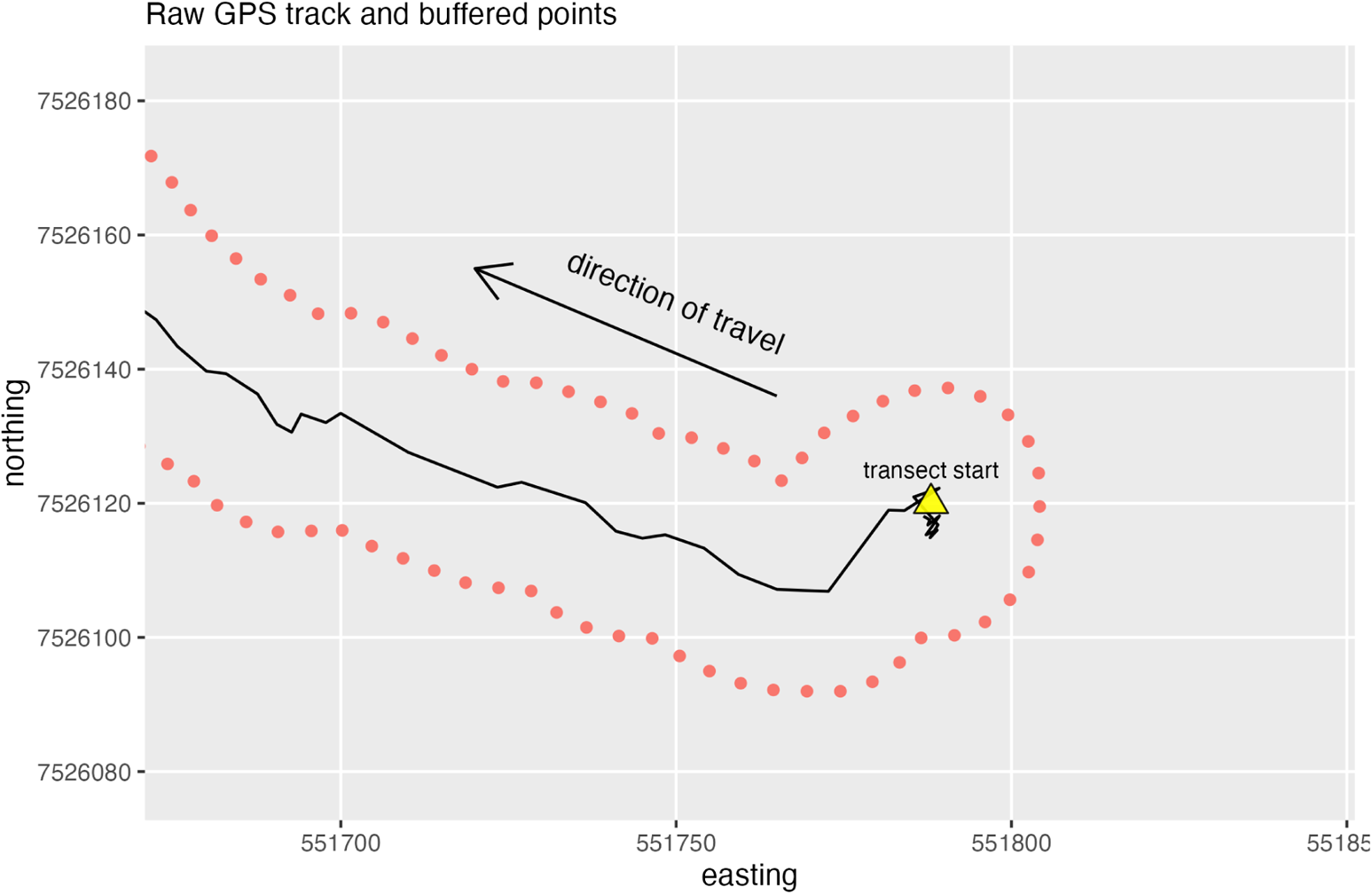
Raw GPS track (black) with points every 5 meters (red) along a 15 meter buffer.

### Manually choose the side of the buffer that will be used as the re-drawn GPS track in QGIS

We identified which of the buffered points were on the left or the right of the raw GPS track. Unfortunately, we were unable to completely automate this step. In QGIS, we manually removed the rounded ends of the buffer points and chose the starting point of the line. Finally, we exported the edited points as a shapefile.

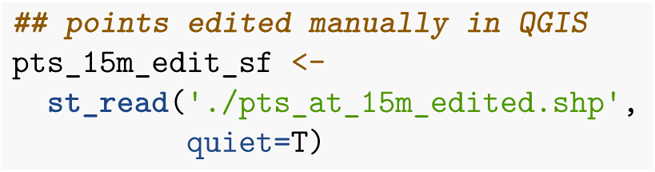

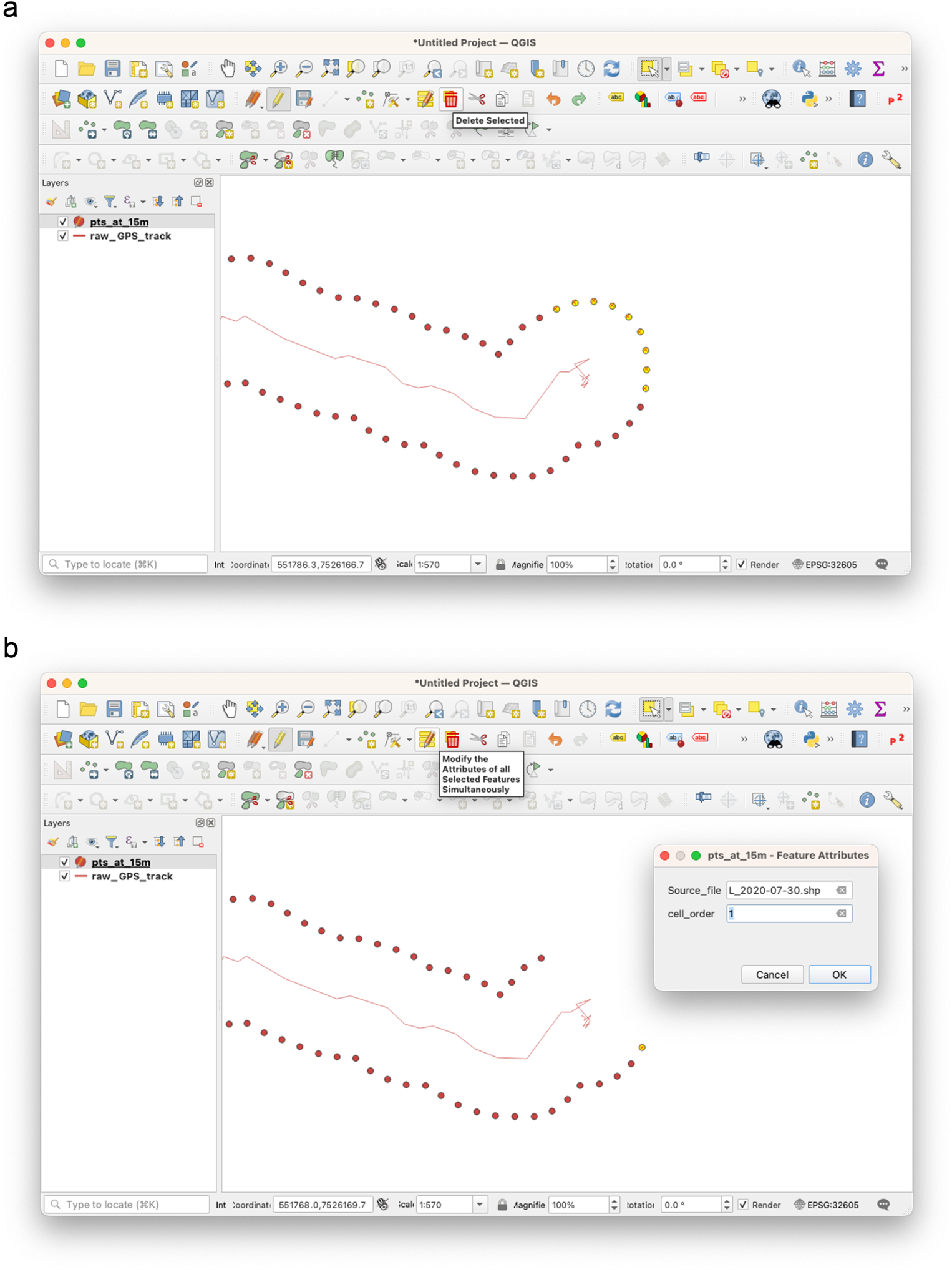
QGIS screenshot: a) remove the buffer points at the start and end of the walk (end not shown). b) we designated the start of the left track by modifying the “cell order” attribute of the first point on the left side of the raw GPS track.

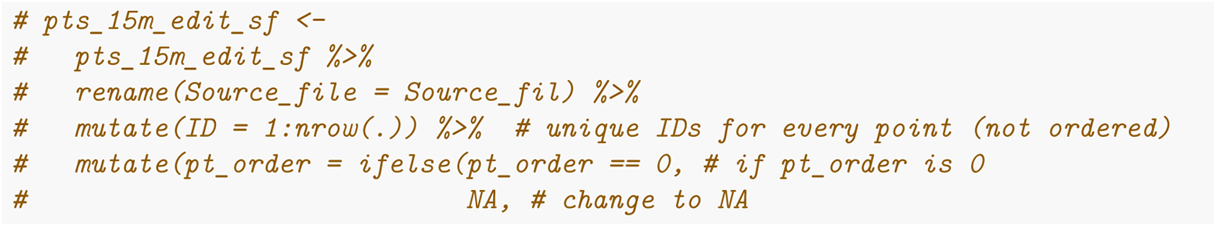
After editing in QGIS, the first buffered point is identified as a 1 in the “pt_order” attribute. All other points are 0 because they have not been assigned a “pt_order” value.

### Order the points from start to end in R

Ordering points gives direction to the path. This is necessary for turning a series of points into a line. We ordered the points using proximity to the hand-selected first point (pt_order = 1). While we were able to automate this step, it takes time for the code to run (about 30 points per second). Fortunately, for just the left side, only half of the points had to be processed (i.e., given about 7000 points in this track, only about 3500 points were processed)

To order the points on one side, we step through “pts_15m_edit_sf” one point at a time using a primary **while()** loop. Starting with the hand-selected first point, we used a nested, secondary **while()** loop, **sf::st_buffer()**, and **sf::st_intersection()** to find the next point within 5 m of the first. If a point was not located within the 5 m buffer, the secondary loop iterates through larger buffer distances. Once the point closest to the first was found, we identified it as “pt_order” = 2. This completed the first iteration of the primary loop. The next iteration of the primary loop repeats these steps to find each successive point. The primary loop was terminated once the secondary loop exceeded a buffer distance 15 m, meaning we had reached the end of the transect.

NOTE for mac users: process multiple transects at a time using multiple cores: https://www.gouskova.com/ 2016/03/09/running-r-on-multiple-cores-mac-os/

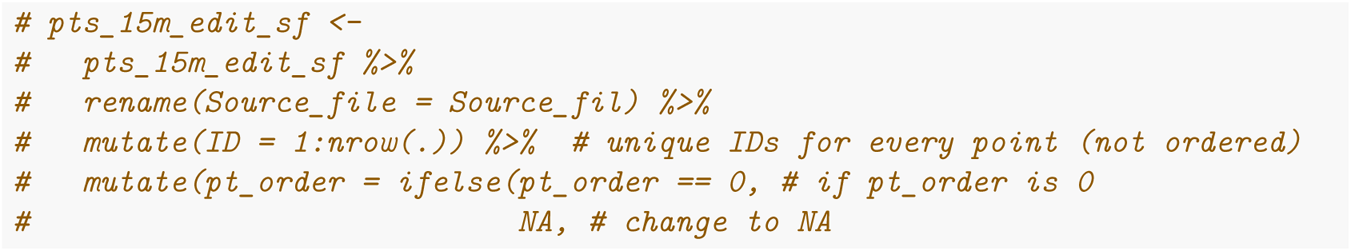

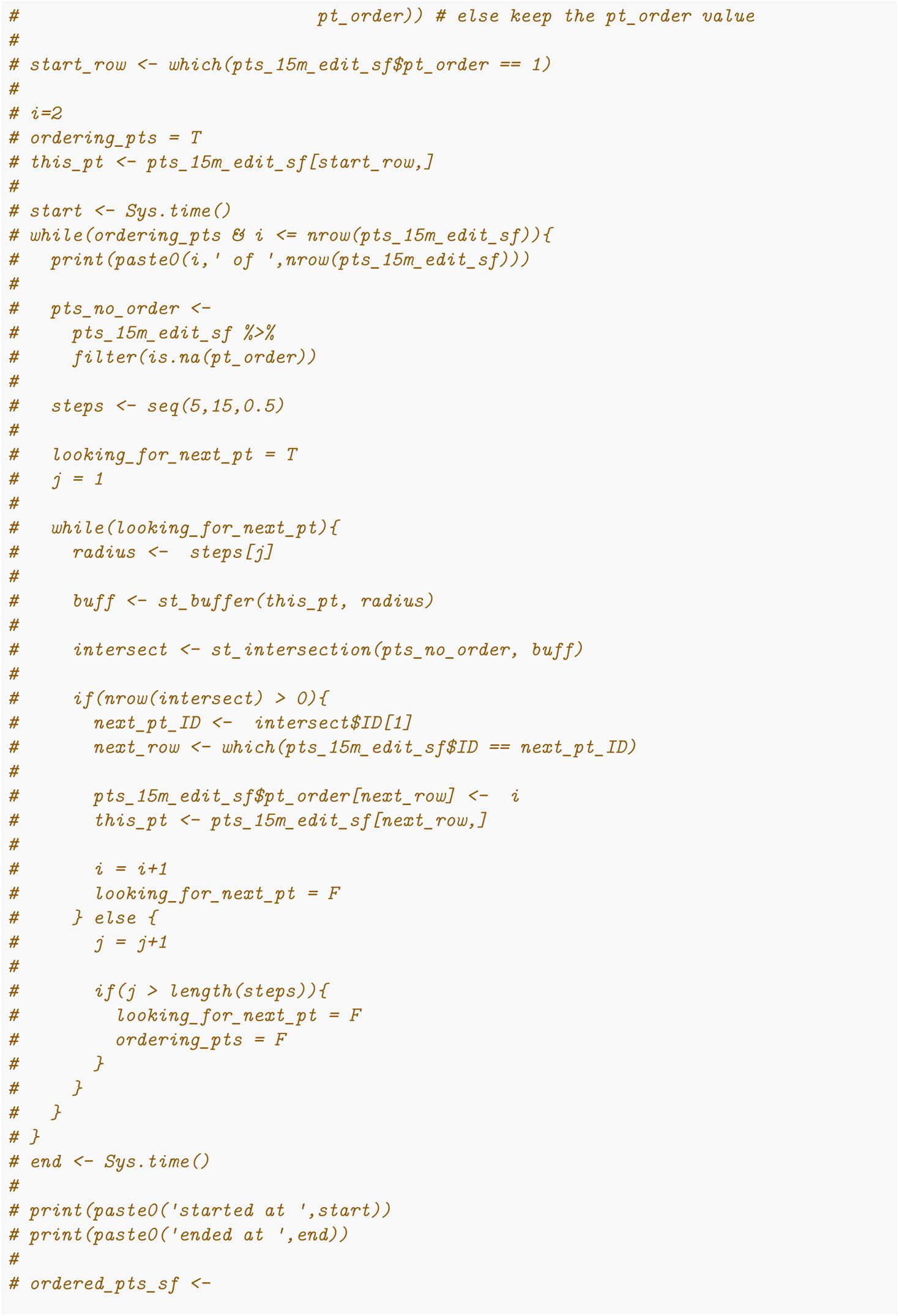

### Completion of the ordered points

There were 3,499 points on the left side of this transect. It took about two minutes to run, or about 29 points per second.

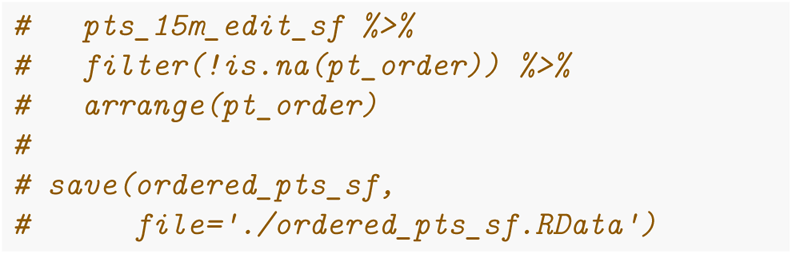

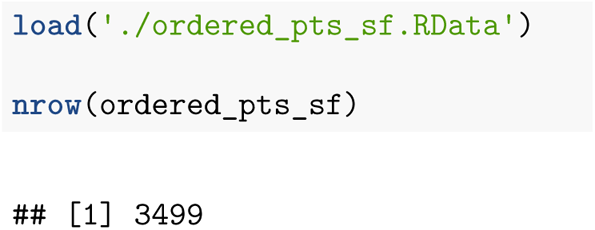
Ordered points starting from the beginning of the transect. Points on the opposing (right) side have been removed.

### Attribute ordered points with changepoint values

We attributed the ordered points with the changepoint classifications. Each changepoint carries a vegetation classification, location, date, time, and unique ID (Fig. Sxx).

#### Read in the changepoints for this transect and prepare a for() loop that iterates through each changepoint

To prepare this loop, we create a vector containing all the unique changepoint IDs: “chpt_ID”. We also created an empty list to fill as the loop runs. Finally, “last_pt_order” is set to one, which is the start of the ordered points.

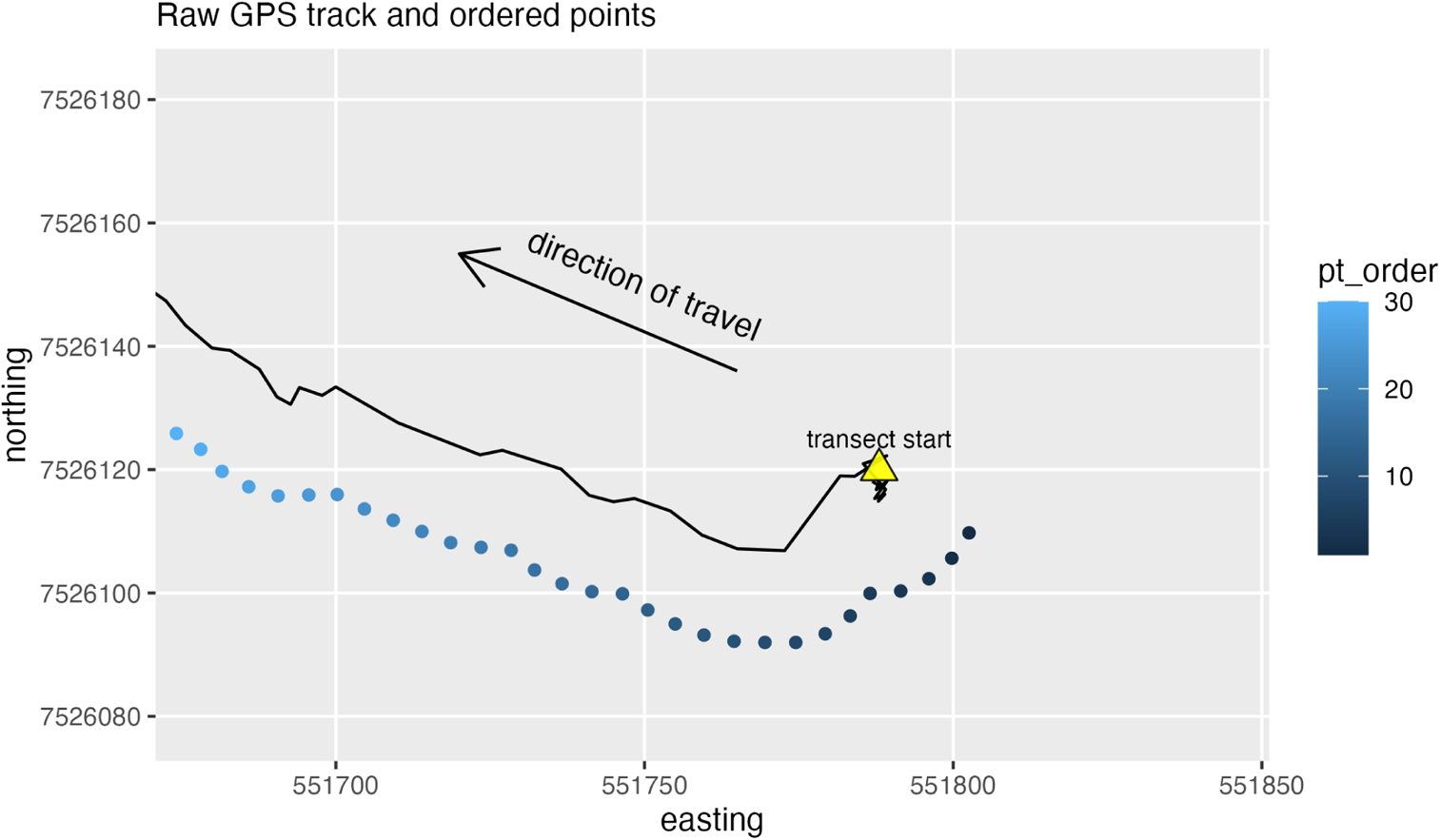

#### Find the ordered point closest to each changepoint

The ordered point closest to each changepoint is the start of that changepoint segment. The following **for()** loop goes through each changepoint and finds the nearest ordered point. The loop progresses through the ordered points in one direction. Each iteration stores the “last_pt_order”, which is used to exclude all ordered points that have passed through previous iterations. The **if()** statement at the beginning of the loop “breaks out” of the loop if the end of the ordered points has been reached. At the end of each iteration the changepoint ID (“chpt_ID”) and the closest point order (“pt_order”) are stored together as a dataframe and added to “changepoint_order_lst”. Finally, “changepoint_order_lst” is collapsed into a dataframe using **do.call()** and **rbind()**.

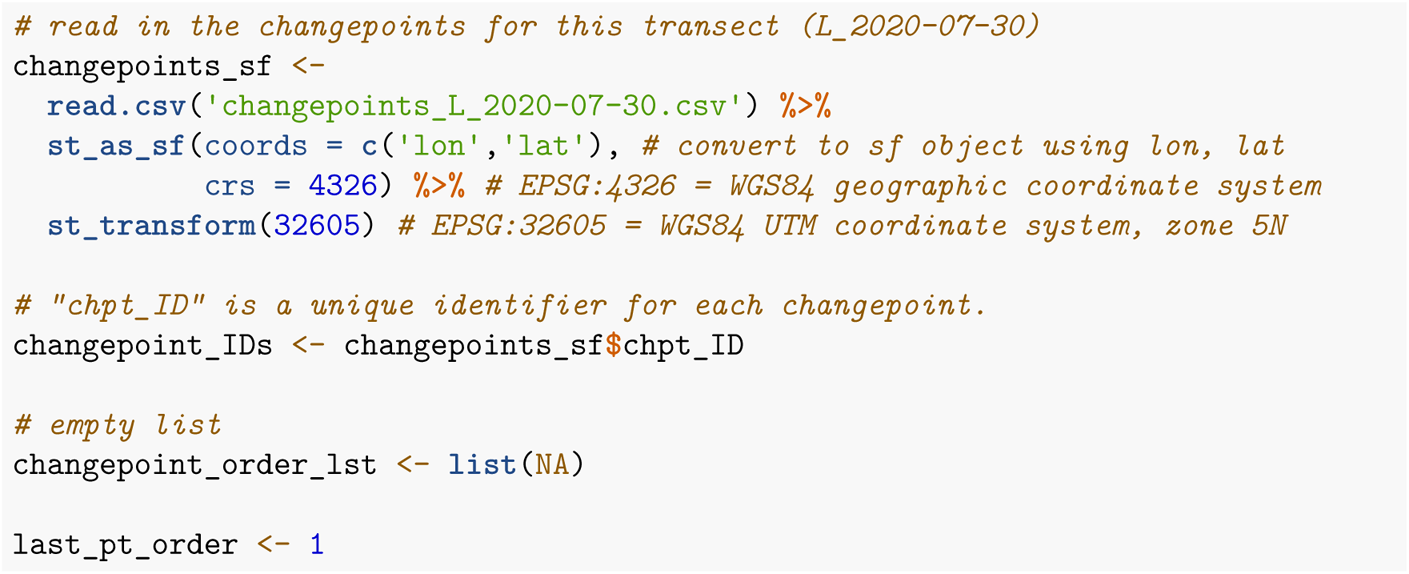

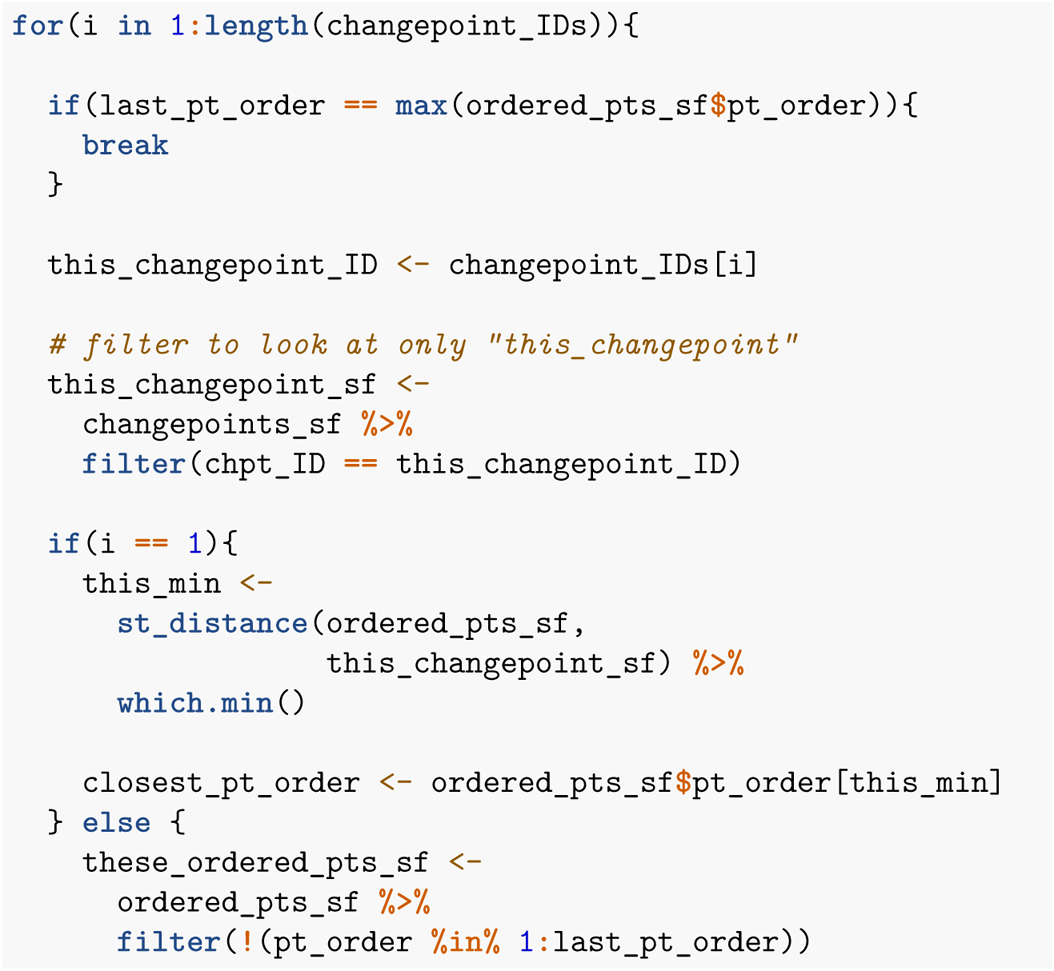

#### Create three vectors that, together, define where each changepoint segment starts and ends along the ordered points

The vectors are: “chpt_ID”, “start” (where the changepoint segment begins along the ordered points), and “end” (where it ends). The starts were determined using the **for()** loop to find closest point order. The ends are the next changepoint’s starting “pt_order” minus 1. The last changepoint ends where the ordered points end.

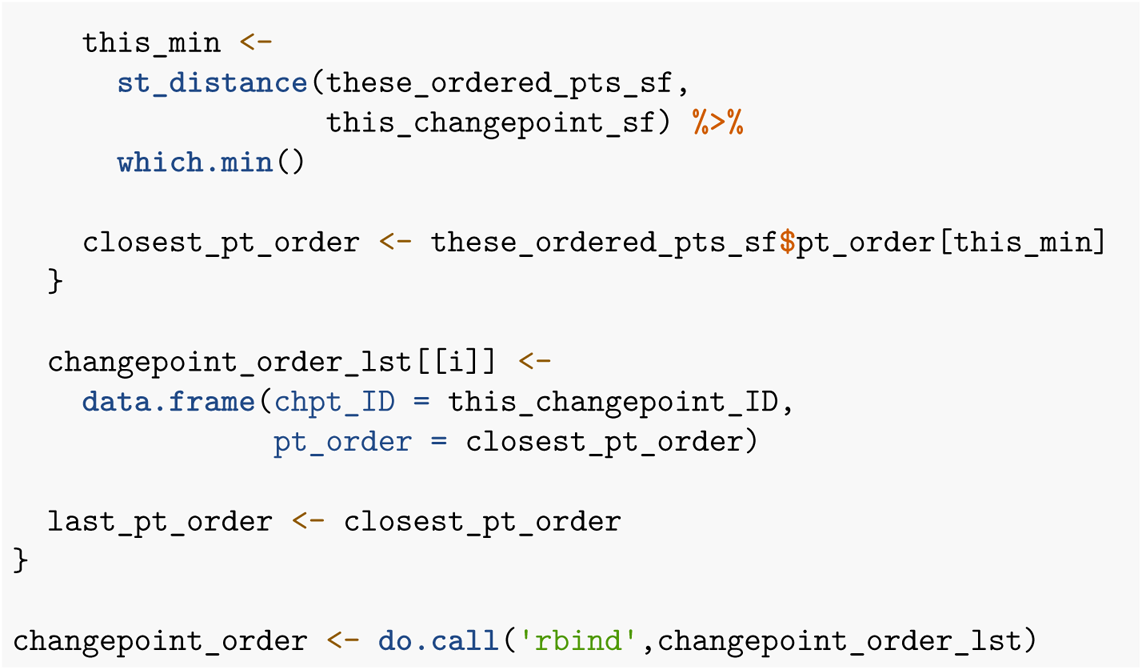

#### Create a dataframe matching every point order to changepoint ID

The function **mapply()** can take multiple input lists and apply a function to them. Here, we use **mapply()** to create a dataframe where each row has the “pt_order” of every point within the changepoint segment. The function creates a dataframe where the “chpt_ID” column has only one value: the changepoint ID. The other column, “pt_order” contain every point along the segment and is created using the colon operator as *start* : *end*. All changepoint segments are collapsed into a dataframe using **do.call()** and **rbind()**. Finally, if there were ordered points before the first changepoint segment (i.e., if the first changepoint was closest to ordered point 12), then all preceding ordered points were given an “NA” as their “chpt_ID”.

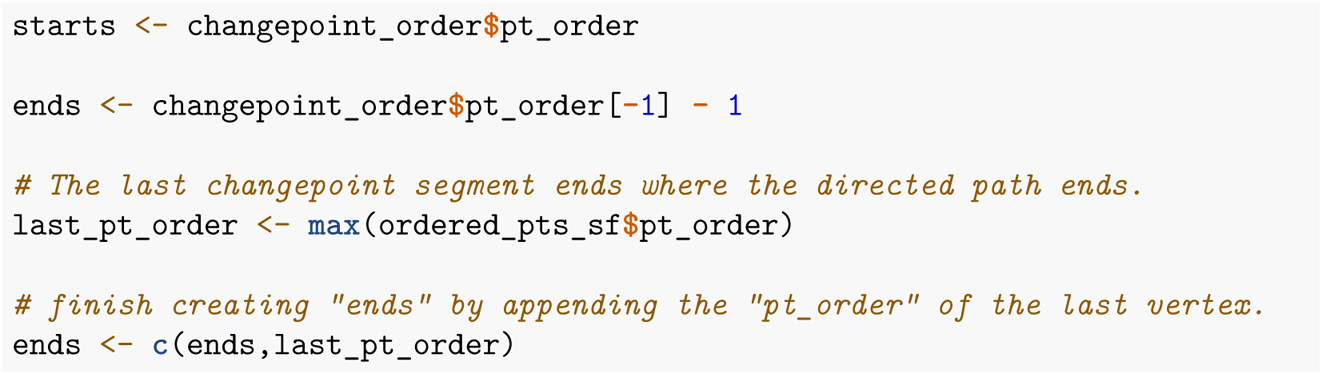

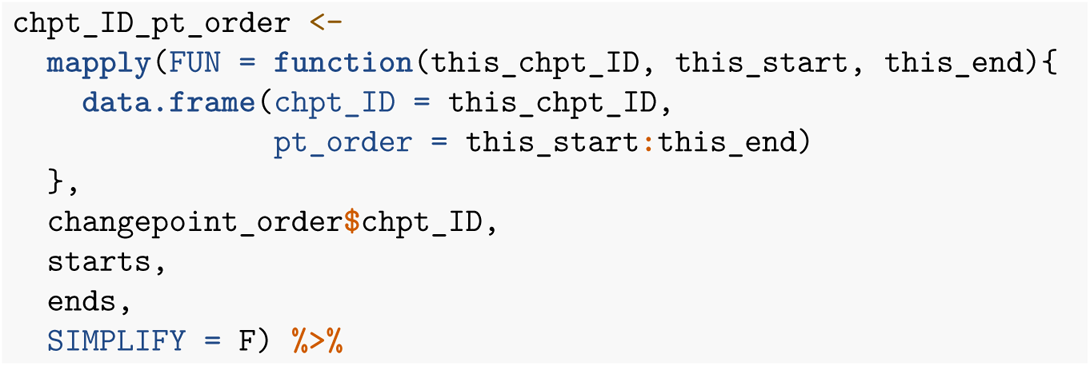

#### Join changepoints to the ordered spatial points

We join the “chpt_ID_pt_order” to the ordered spatial points (“ordered_pts_sf”) using **dplyr::left_join()** by “pt_order.”

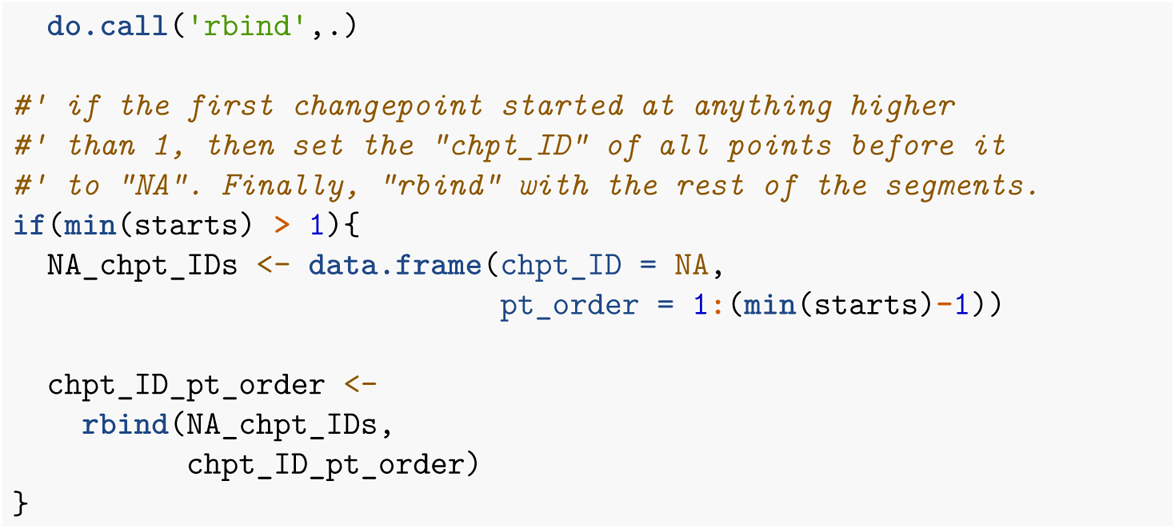

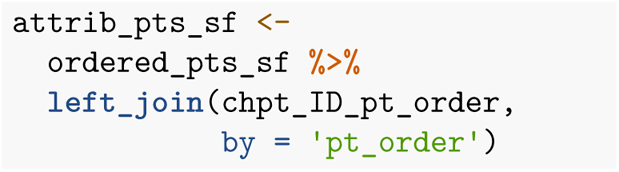
Attributed points for the first two changepoints. The gray points near the start of the directed walk are “NAs” because no changepoint was collected for the first 50 meters of the transect.

### Converting points to lines

We want to convert each spatial point into a line segment that terminates at the next point. However, spatial points alone cannot be converted into a line simply by “connecting the dots”. While we know each spatial point is the “start” of its own segment and the “end” is the next point, the “end” must be defined in a way that maintains the data associated with the spatial point at the start of the segment. Similar to the changepoint segments, these next lines of code create starts and ends of segments, where each segment is attributed by the spatial point at its start.

#### Establish starts and ends of line segments along the attributed points

First, for easier handling later on, we combine ‘x’ and ‘y’ coordinates into singular variables for starts and ends. The starts are the coordinates of each point *i*, and the ends are there coordinates of the next point (*i* + 1). Because the last point has no end, it is removed.

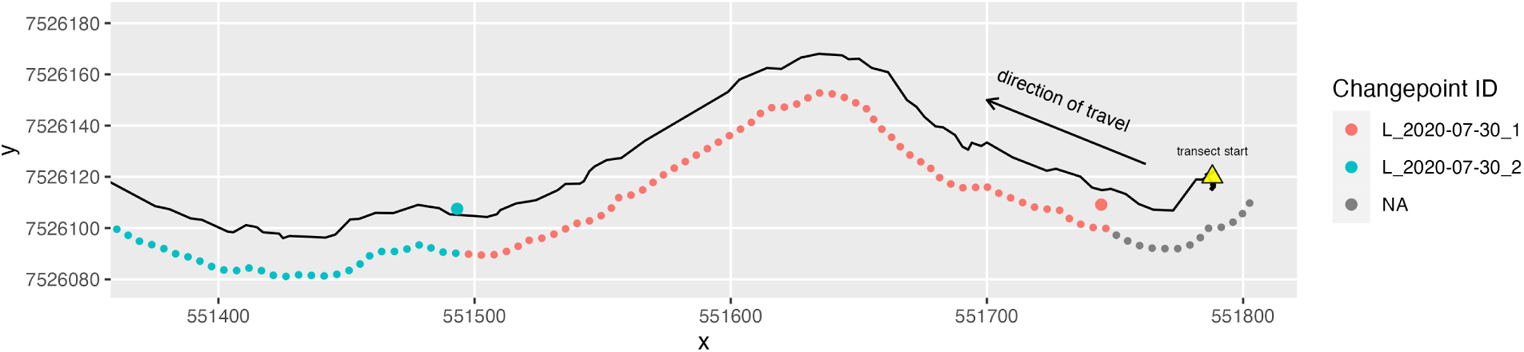

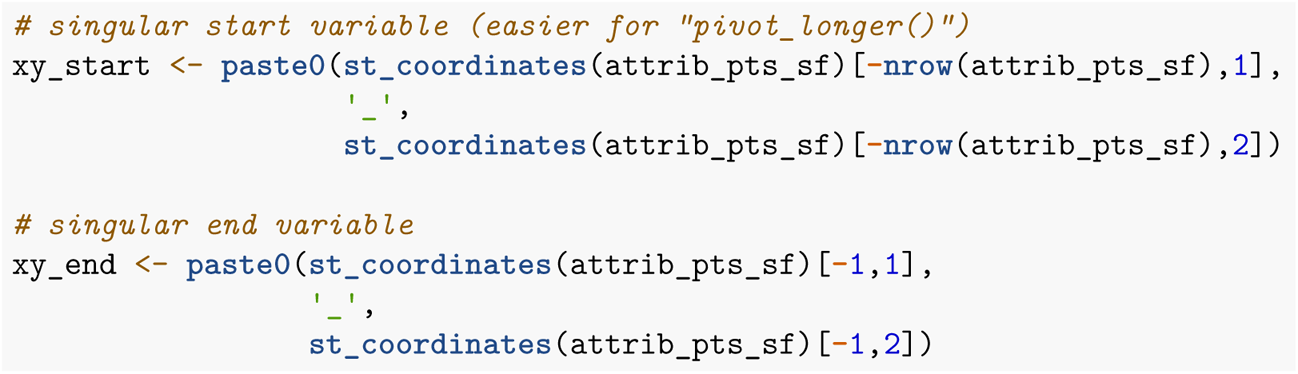

#### Add coordinates to the dataframe as variables

First, we remove the last point using **dplyr::slice()**, then add the start and end coordinates using **dplyr::mutate()**. Lastly, we remove the geometry using **sf::st_drop_geometry()**. Each row of the resulting dataframe represents a line segment but is not a spatial line.

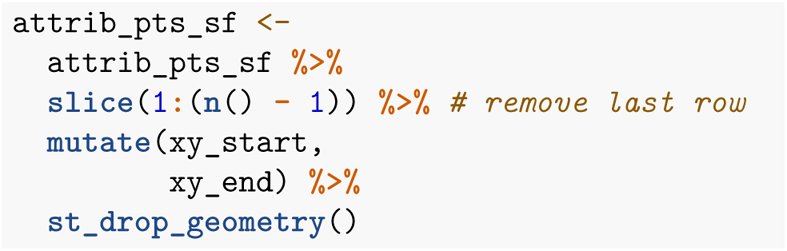

#### Split each row into a pair of two points rows where one is the start of the segment and the other is the end

Using **dplyr::pivot_longer()** we separate each row start and end using the coordinates defined above. We also separate the combined “x” and “y” coordinates into their own columns and convert back to spatial points using **sf::st_as_sf()**. The

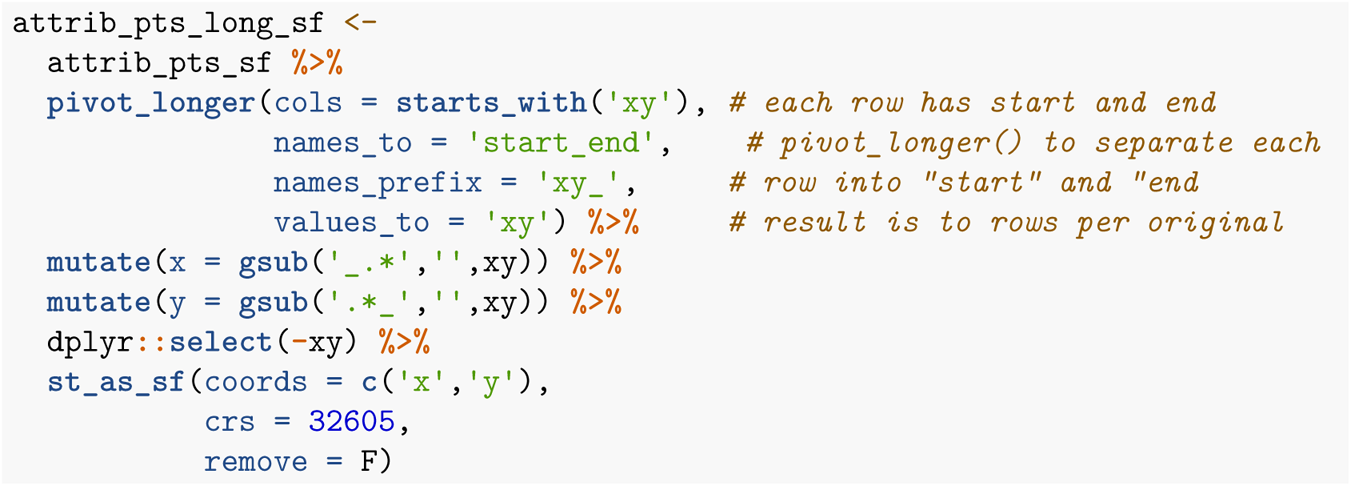

#### Convert paired points to lines

Now, the column “pt_order” acts as a unique ID for each segment of the directed path. This unique ID is shared between each start-end pair of spatial points. We use **dplyr::group_by()** on “chpt_ID” and “pt_order” and **dplyr::summarize()** (summarise function is irrelevant, we use sample size) to collapse each pair into a “MULTIPOINT”. Finally, with **sf::st_cast()** we convert each “MULTIPOINT” into a “LINESTRING” or a spatial line segment where each segment can have unique attributes.

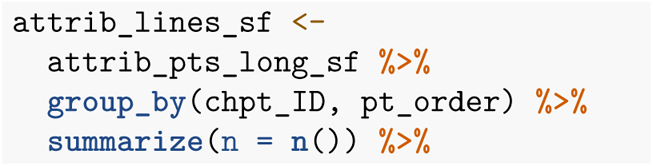

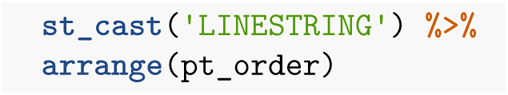

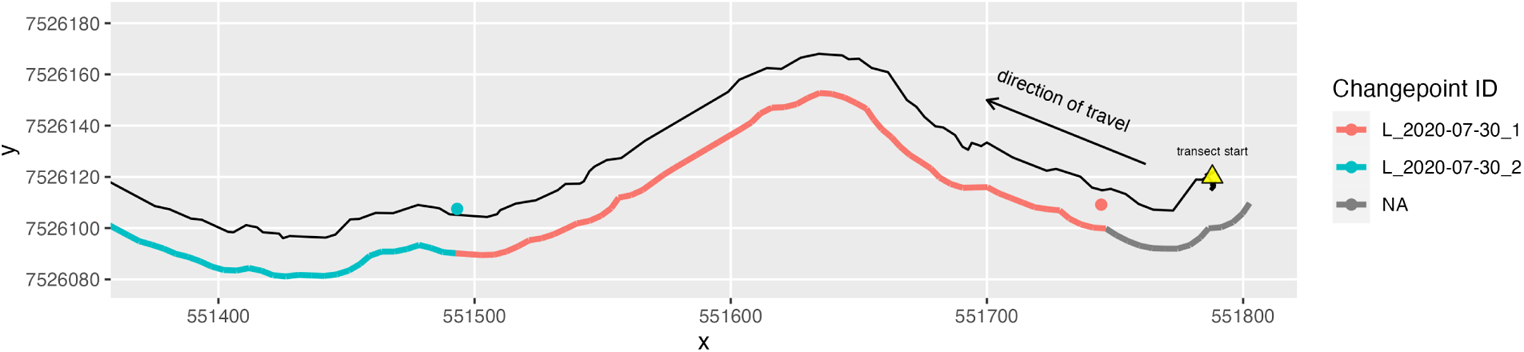
The black line is the original directed walk recorded in the field. The arrow indicates the direction of travel from right to left. The colored line next to the directed walk is the parallel path that we re-drew 15 meters to the left of the walk. The blue and red circles are changepoints. The colors of the path and circles correspond to changepoint classifications that have been attributed to the path. The gray portion of the line is classified as “NA” because no changepoint was collected until shortly after the directed walk began.

## Supplementary Text

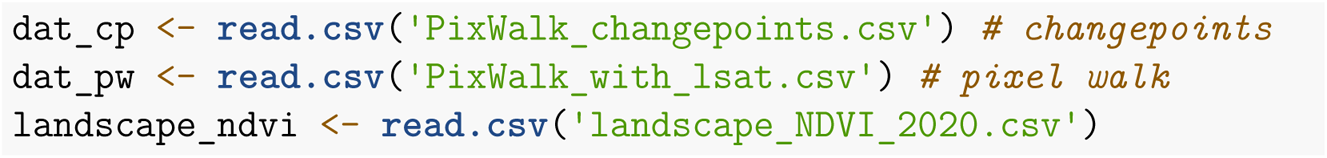

### What elevations were sampled?

Our transects sampled from 250-1300 m asl with a mean of about 700 m asl.

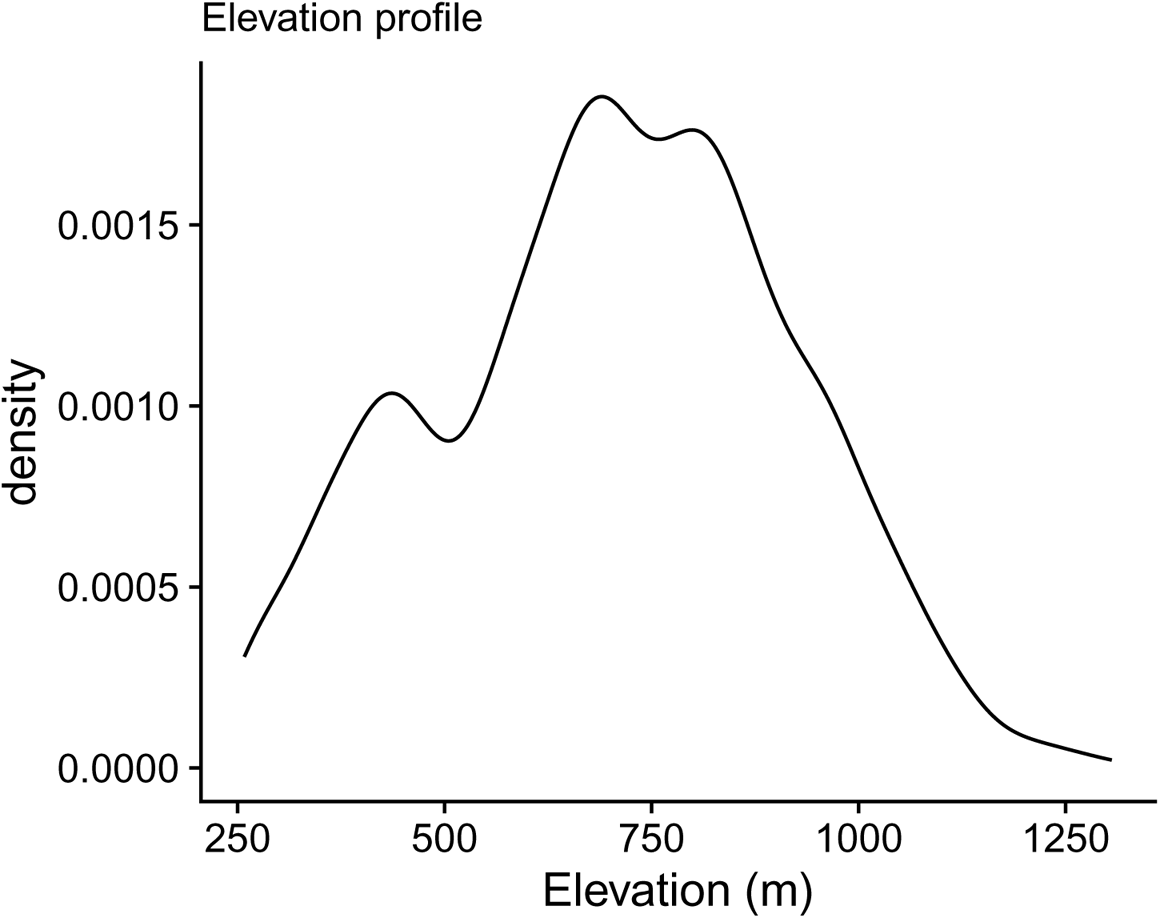

### How many pixels were sampled?

23,213 pixels were sampled for max-greenness out of 24,395 total pixels. The remaining pixels did not have max-greenness estimates due to poor fits of phenological curves during the 2020 growing season.

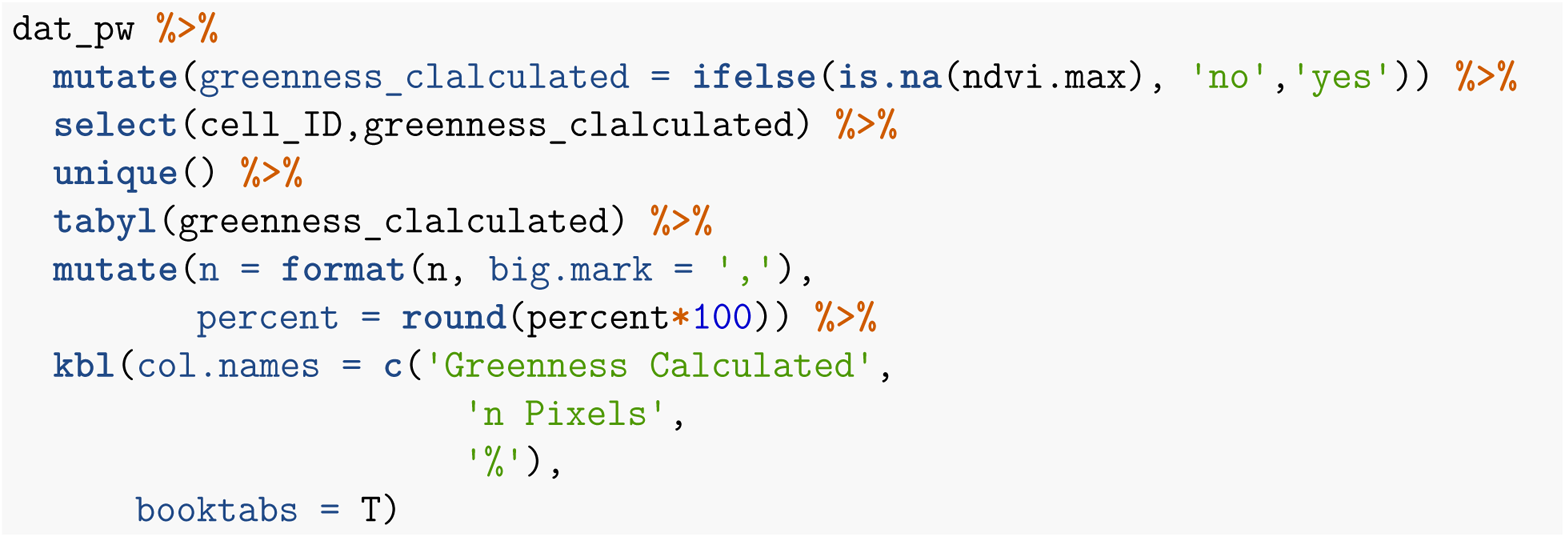

| Greenness Calculated | n Pixels | % |
| --- | --- | --- |
| no | 1,182 | 5 |
| yes | 23,213 | 95 |

### How many pixels were sampled per day?

We sampled 90 to 2,095 pixels sampled each day with an average of 1,220 per day.

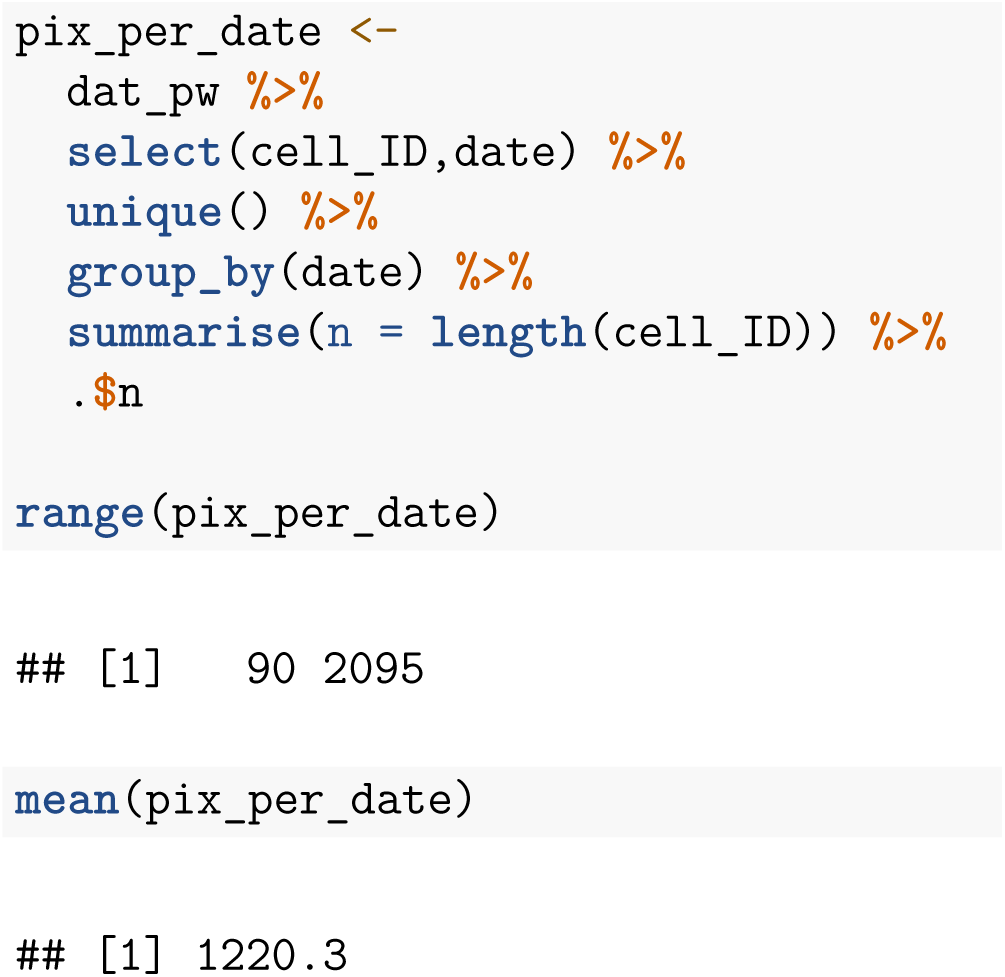

### What was the total length of all transects?

We collected 639 km of vegetation transects.

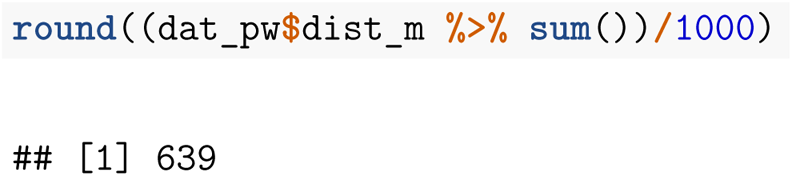

### How much of each pixel was sampled with our transects?

Transects could have intersected Landsat pixels in various ways. This shows that most often transects were capturing close to 30 m of Landsat pixels.

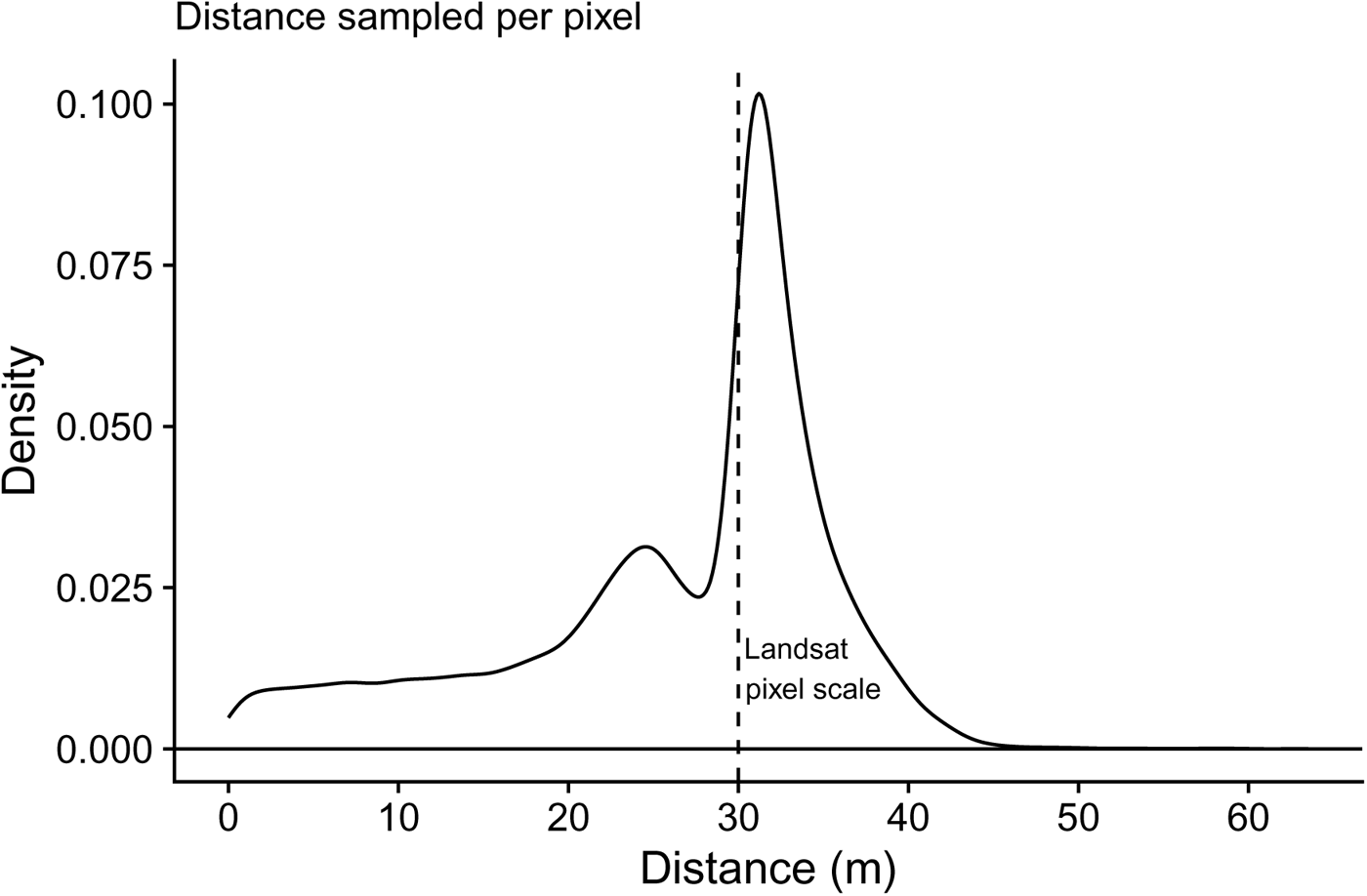
Sum of transect lengths for each pixel. Vertical dashed line is the resolution of a square Landsat pixel (30 m)

### Multiple regression

#### Pixel proportions

To set up the regression, we need a dataframe of all pixels and veg proportions. Each row of the matrix is a pixel. The first column is the pixel value, maximum NDVI. All other columns are the observed covrPFTs and their values are the proportions. The table below gives the first five rows and five covrPFTs of an example dataframe.

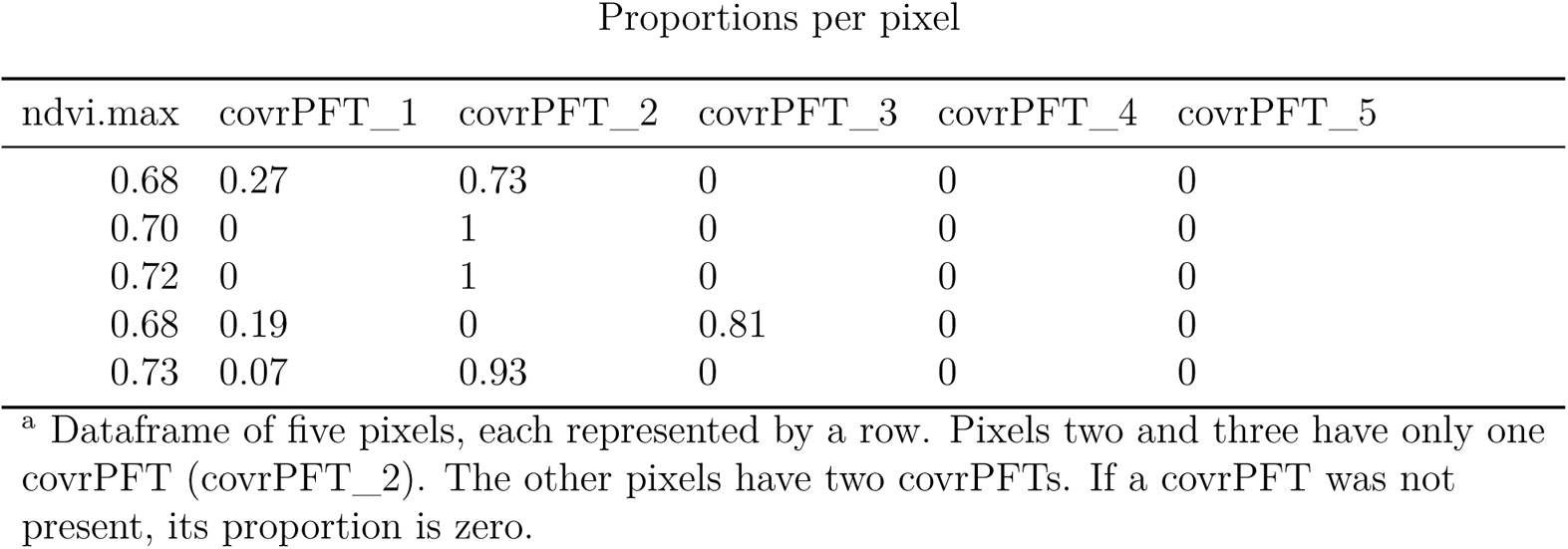

Proportions per pixel
| ndvi.max | covrPFT_1 | covrPFT_2 | covrPFT_3 | covrPFT_4 | covrPFT_5 |
| --- | --- | --- | --- | --- | --- |
| 0.68 | 0.27 | 0.73 | 0 | 0 | 0 |
| 0.70 | 0 | 1 | 0 | 0 | 0 |
| 0.72 | 0 | 1 | 0 | 0 | 0 |
| 0.68 | 0.19 | 0 | 0.81 | 0 | 0 |
| 0.73 | 0.07 | 0.93 | 0 | 0 | 0 |

<sup>a</sup> Dataframe of five pixels, each represented by a row. Pixels two and three have only one covrPFT (covrPFT\_2). The other pixels have two covrPFTs. If a covrPFT was not present, its proportion is zero.

#### Fitting linear model

The dataframe of proportions is the input for the linear model. The dependent variable is the pixel value, “ndvi.max”, and the independent variables are the proportions from all columns, given by “.”, with no intercept, given by “-1”.

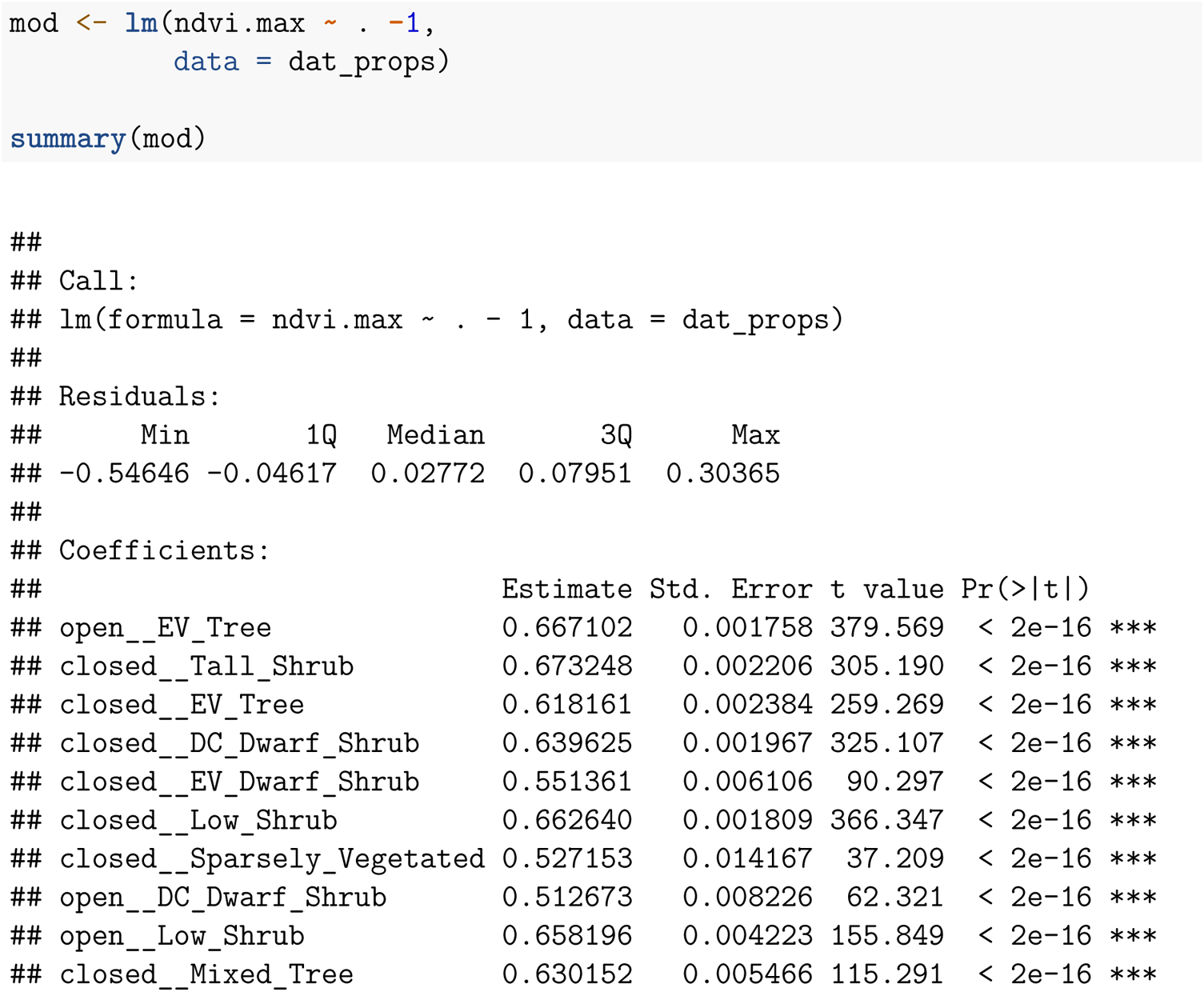

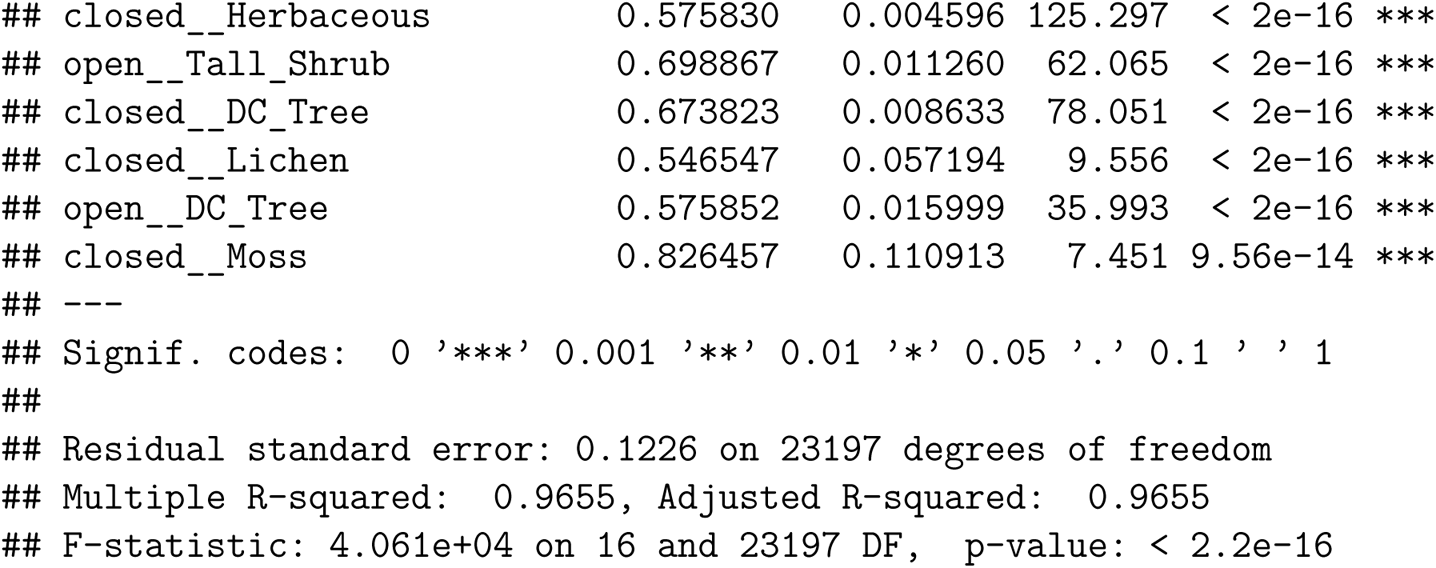

We are primarily interested in the “Estimate” and “Std. Error” for estimating mean NDVI for each vegetation class. Due to large sample size, all estimates have large t- and p-values. That said, it is expected that NDVI estimates are significantly different than zero.

### Homogenous Pixels

Some pixels were classified with only one combination of overstory and understory vegetation. We analyzed these separately to validate the inclusion of heterogenous pixels.

#### How many pixels (with maximum NDVI) had only one cPFT?

In the main text, we don’t report all cPFTs. The following calculates the number of pixels of *all* cPFTs. To maintain consistency

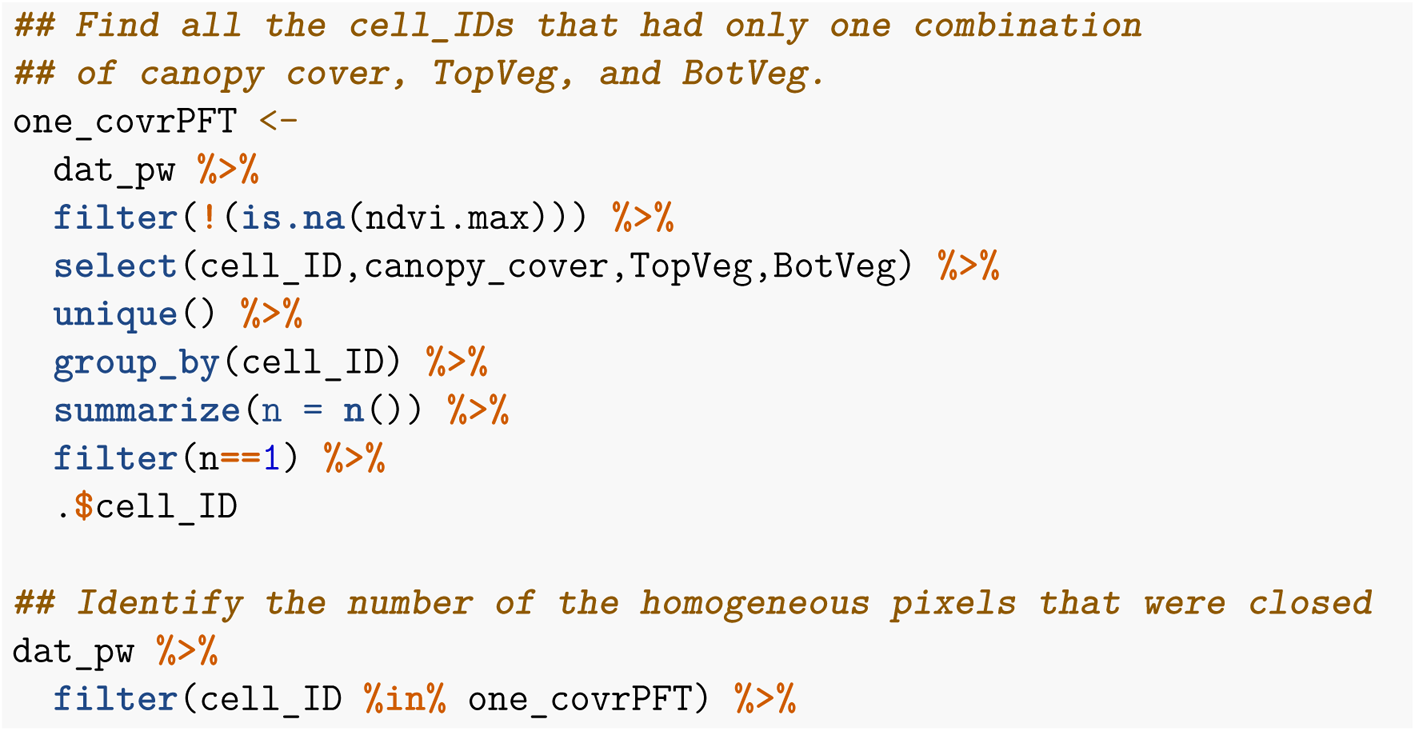

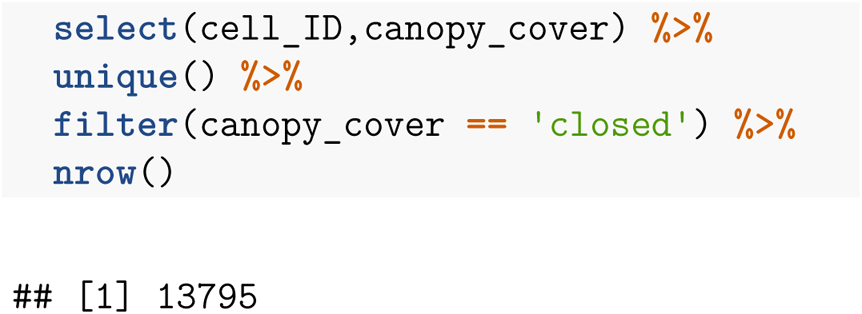

13,795 pixels had only one closed overstory PFT with one understory PFT. In the main text we report only the covrPFTs with >1km sample length. The excluded covrPFTs account for Figure 2 reporting five less pixels than above.

#### Using the the homogeneous cPFT pixels, what is the mean greenness for each of the cPFTs?

We calculated arithmetic pixel means using the sum of their NDVI values divided by their sample size. The cPFTs that were not reported in the main text are *italicized*.

| cPFT | mean_greenness | n_pixels |
| --- | --- | --- |
| *Moss* | *0.77* | *1* |
| Deciduous_Tree | 0.68 | 185 |
| Tall_Shrub | 0.67 | 2,612 |
| Low_Shrub | 0.66 | 3,856 |
| Deciduous_Dwarf_Shrub | 0.64 | 3,326 |
| Mixed_Tree | 0.63 | 455 |
| Evergreen_Tree | 0.62 | 2,340 |
| Herbaceous | 0.57 | 614 |
| Evergreen_Dwarf_Shrub | 0.55 | 334 |
| *Lichen* | *0.54* | *4* |
| Sparsely_Vegetated | 0.53 | 68 |

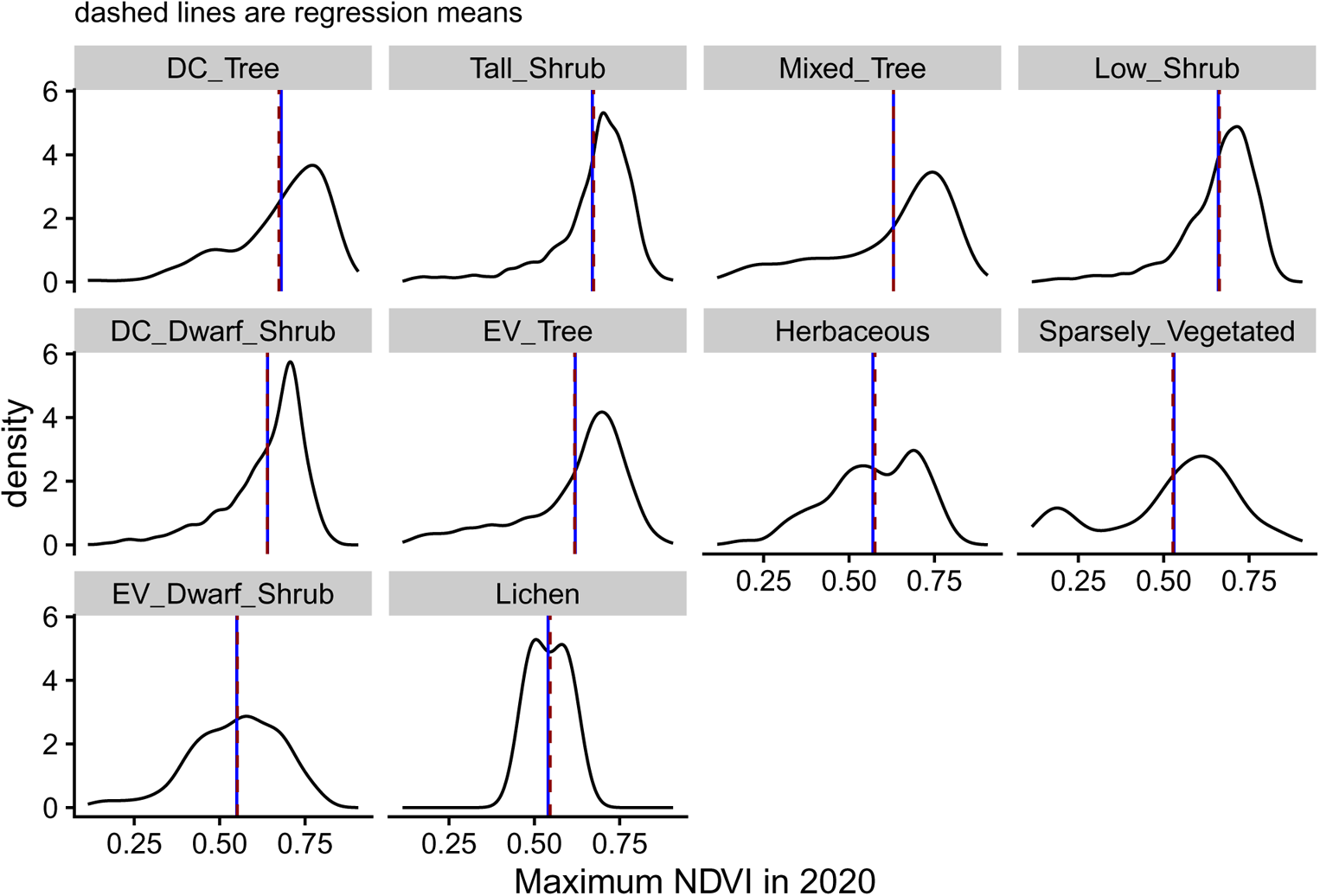

#### How many pixels (with maximum NDVI) had only one oPFT?

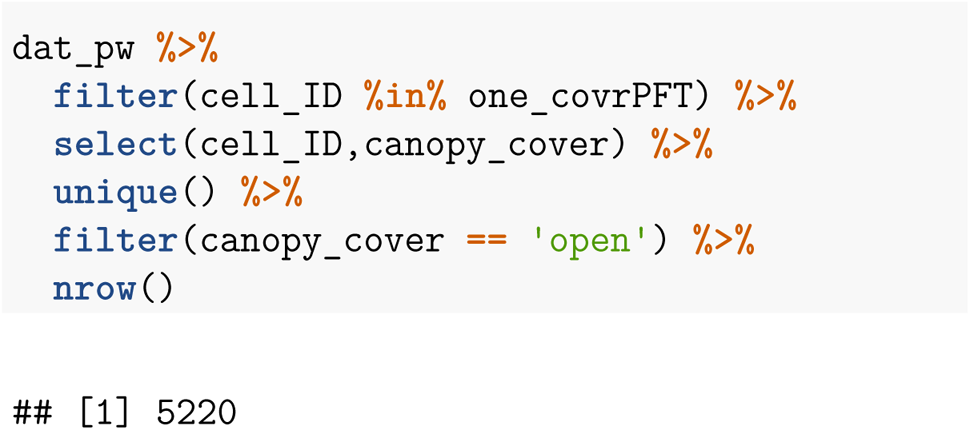

5,438 pixels had only one open overstory PFT with one understory PFT.

#### Using the the homogeneous oPFT pixels, what is the mean greenness for each of the oPFTs?

Arithmetic pixel means:

| oPFT | mean_greenness | n_pixels |
| --- | --- | --- |
| Tall_Shrub | 0.70 | 95 |
| Evergreen_Tree | 0.67 | 4,129 |
| Low_Shrub | 0.66 | 747 |
| Deciduous_Tree | 0.57 | 55 |
| Deciduous_Dwarf_Shrub | 0.51 | 194 |

#### Regression of homogenous pixels

To further validate the regression method, we repeated multiple regression for the same pixels used to calculate arithmetic mean greenness (i.e., homogenous pixels). The means were exactly the same. Error was mostly similar when low, but regression error grew exponentially as arithmetic error grew linearly.

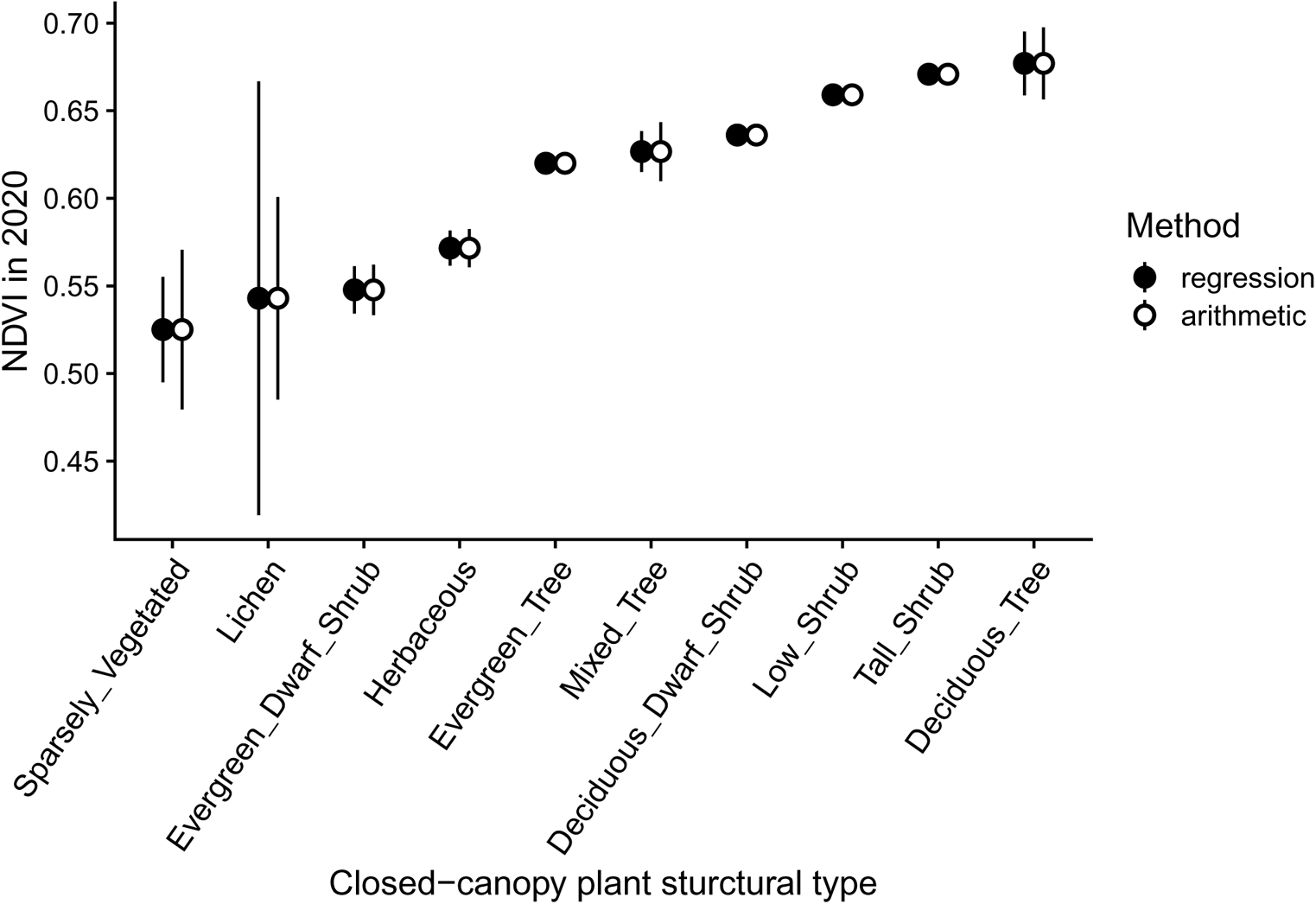

#### What PFTs are found in the understory?

Satellite observed greenness is likely influenced by understory vegetation. The following tables list the understory PFTs found underneath all of the oPFTs and the same cPFTs.

| overstory_oPFT | understory_PFT | dist_km | percent |
| --- | --- | --- | --- |
| Tall_Shrub | Deciduous_Dwarf_Shrub | 2.0 | 57 |
|  | Evergreen_Dwarf_Shrub | 0.5 | 14 |
|  | Moss | 0.4 | 13 |
|  | Herbaceous | 0.3 | 9 |
|  | Lichen | 0.3 | 8 |
| Evergreen_Tree | Deciduous_Dwarf_Shrub | 38.0 | 29 |
|  | Herbaceous | 35.4 | 27 |
|  | Lichen | 26.0 | 20 |
|  | Evergreen_Dwarf_Shrub | 23.7 | 18 |
|  | Tussock | 5.8 | 4 |
|  | Moss | 3.7 | 3 |
|  | Low_Shrub | 0.0 | 0 |
| Low_Shrub | Deciduous_Dwarf_Shrub | 18.3 | 76 |
|  | Evergreen_Dwarf_Shrub | 3.5 | 15 |
|  | Lichen | 1.0 | 4 |
|  | Moss | 1.0 | 4 |
|  | Herbaceous | 0.3 | 1 |
| Deciduous_Tree | Evergreen_Dwarf_Shrub | 1.5 | 85 |
|  | Deciduous_Dwarf_Shrub | 0.2 | 10 |
|  | Herbaceous | 0.1 | 5 |
| Deciduous_Dwarf_Shrub | Evergreen_Dwarf_Shrub | 4.6 | 71 |
|  | Moss | 1.1 | 17 |
|  | Deciduous_Dwarf_Shrub | 0.7 | 11 |
|  | Lichen | 0.1 | 1 |

<sup>a</sup> All understory PFTs observed under each oPFT. Distance (km) is the sum of transect segments with the observed combination of oPFT and understory PFT. Percent is the proportion of each understory PFT relative to the total length of the overstory oPFT.

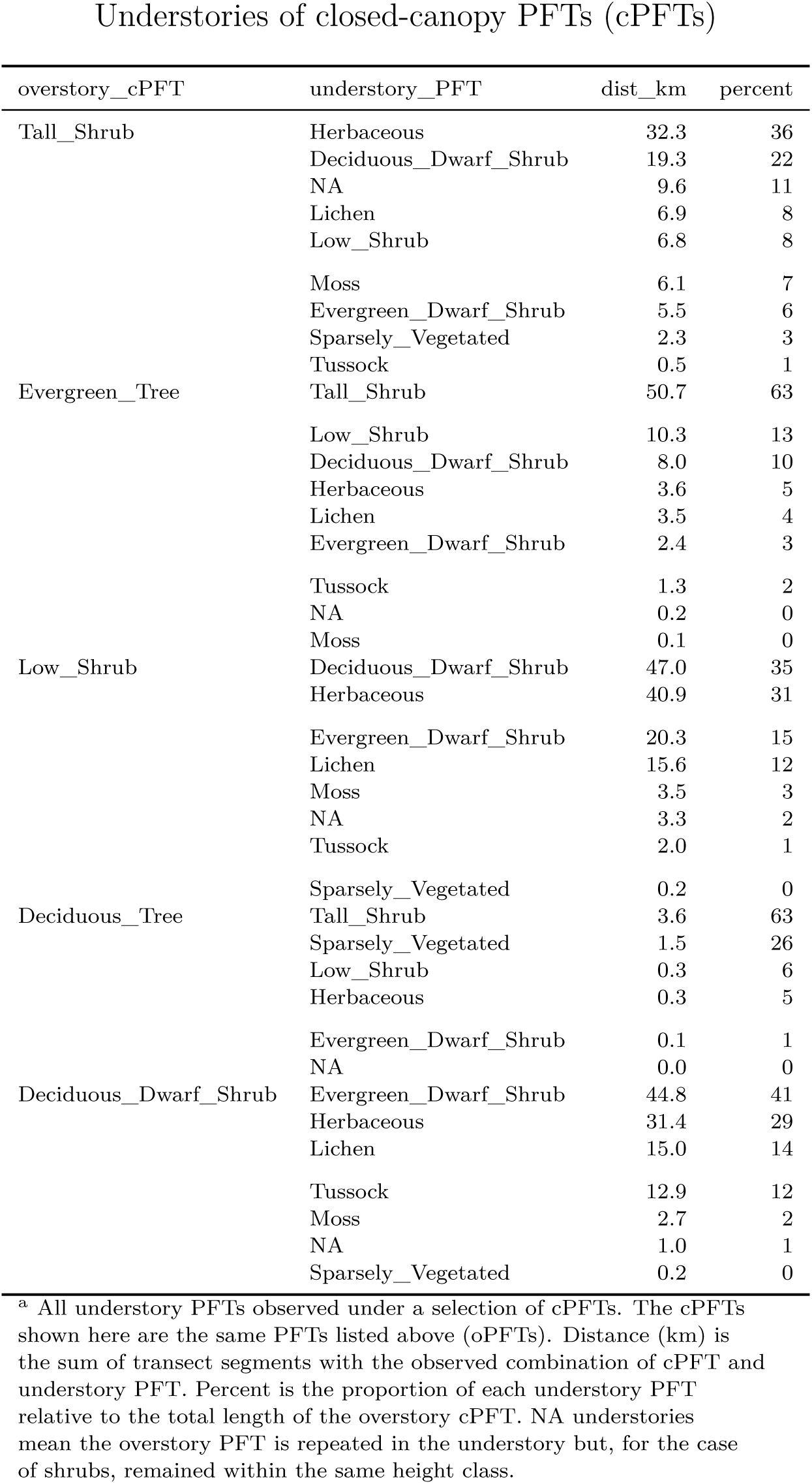

